# Global and local origins of trial-to-trial spike count variability in visual cortex

**DOI:** 10.1101/2025.08.08.669442

**Authors:** Anna J. Li, Ziyu Lu, Alexander E. Ladd, Pascha Matveev, Eric Shea-Brown, Nicholas A. Steinmetz

**Author notes:** Authors contributed equally (Anna J. Li, Ziyu Lu). Authors jointly supervised (Eric Shea-Brown, Nicholas A. Steinmetz).

## Abstract

Sensory neuron spiking responses vary across repeated presentations of the same stimuli, but whether this trial-to-trial variability represents noise versus unidentified signals remains unresolved. Some of the variability can be attributed to correlations between neural activity and arousal, locomotion, and other overt movements. We hypothesized that correlations with global activity factors, i.e., patterns of neural activity observable in other brain regions, may explain additional variability in spike count responses of visual cortical neurons. To test this, we used Neuropixels 2.0 probes to record neural activity in mouse primary visual cortex (V1) while subjects passively viewed images. We recorded videos of behavior alongside neural activity from other brain regions, either spiking activity of neural populations in anterior cingulate area (ACA) or widefield calcium signals from across the dorsal cortex. We then used a model based on reduced rank regression to partition the explainable variability of visual cortical responses by source. Some of the trial-to-trial variability in V1 spike counts was attributable to locally shared patterns of activity uncorrelated with either behavior or global activity patterns. Locally shared activity patterns explained trial-to-trial variability that was in excess of Poisson spike generation. Of the parts of variability attributable to non-local sources, global cortical activity predicted significantly more V1 spike count variability than behavioral factors. Additionally, behavioral factors explained little variability uniquely and comprised a geometric subspace of the globally predictable V1 activity. Finally, optogenetically perturbing ACA directly impacted V1 activity, and ACA activity patterns predicted V1 spike count variability even on trials without overt behaviors. Our data indicate that globally shared factors from other cortical areas contribute substantially to shared spike count variability in V1, with only a minority of shared variability confined to local V1 circuits.

## Introduction

Spike trains produced by sensory cortical neurons in response to repeated presentations of identical stimuli can vary substantially from trial to trial (Bair and Koch, 1996; Tolhurst et al., 1983). This variability poses challenges for theories of neural computation, which must address how effective information processing occurs despite unreliable neuronal signaling. While spike generation within a neuron can be highly reliable (Mainen and Sejnowski, 1995), it remains unclear what proportion of variability is attributable to processes at the single-neuron versus network level. For example, spiking variability may arise via irreducible sources such as stochastic quantal release from synapses (Fatt and Katz, 1952) or other biophysical noise sources (Faisal et al., 2008). Additionally, local network mechanisms may give rise to shared chaotic fluctuations (Huang et al., 2019; van Vreeswijk and Sompolinsky, 1996) or other types of shared local fluctuations (Ahmadian and Miller, 2021; Renart et al., 2010), which may drive spiking variability (Pattadkal et al., 2024). Accordingly, neurons in cortical networks have been modeled as Poisson spike generators, with super-Poisson components of variability shared at the population level (Goris et al., 2014). Shared, locally-correlated fluctuations can constrain the nature and precision of encoded information (Averbeck et al., 2006; Rumyantsev et al., 2020; Stringer et al., 2019a; Zylberberg et al., 2017). Thus, a wide variety of different mechanisms may drive neural variability, but determining which mechanisms dominate in neural circuits remains an empirical and theoretical challenge. In particular, investigations of these phenomena typically focus on single regions, and whether the mechanisms underlying locally shared fluctuations extend beyond individual cortical regions to broader distributed networks remains unclear.

A key question in understanding cortical variability is the relative contribution of local circuit dynamics versus global network influences. If most variability patterns are confined within cortical areas, this would support models of cortical function that emphasize independent computational units composed of local recurrent connectivity. Conversely, if variability patterns largely reflect coordinated, global activity across brain regions, this would suggest that cortical areas operate as components of distributed networks. The resolution of this question has implications for both the mechanistic basis of cortical variability and the organizational principles of cortical computation.

A salient feature governing shared fluctuations within larger scale networks is that only a subset of signals within a cortical region may be transmitted to neurons outside that region, with the rest residing in so-called “null spaces” in which activity does not influence target neurons (Churchland and Shenoy, 2024; Kaufman et al., 2014). The “potent space” activity, or multi-dimensional correlations between regions, are often termed “communication subspaces” and have been measured in a variety of brain regions and behavioral contexts (Ebrahimi et al., 2022; MacDowell et al., 2023; Semedo et al., 2019; Srinath et al., 2021; Steinmetz et al., 2019; Veuthey et al., 2020). However, despite recent advances in accounting for multi-region correlations at the population level (Akella et al., 2025; Huang et al., 2019), the impact of shared fluctuations across regions on trial-to-trial spike count variability in single neurons of sensory cortex is unclear.

Some of the cortical activity that is shared between neurons and across regions is attributable to spontaneous overt behavior. Specifically, measurable behaviors of subjects (e.g. from videos) contain predictive information about activity fluctuations in both sensory cortex and more broadly (Manley et al., 2024; Musall et al., 2019; Steinmetz et al., 2019; Stringer et al., 2019b; Yin et al., 2025). Though these behavior-related activity fluctuations are widespread, and may be species-dependent (Liska et al., 2024; Talluri et al., 2023), they nevertheless account for only a subset of the activity patterns shared across regions. While it is known that some activity in sensory cortex is therefore correlated with these globally-shared patterns, prior measurements of this property were primarily made with calcium imaging, precluding the quantification of the extent to which trial-to-trial spike count variability is attributable to these factors. We hypothesized that globally-shared cortical activity patterns may be a major source of trial-to-trial spike count variability, and overlap with or even subsume the variability attributable to behavior.

Here, we recorded spikes from hundreds of neurons in mouse primary visual cortex (V1) with four-shank Neuropixels 2.0 probes (Steinmetz et al., 2021). We employed a method not previously used for partitioning neuronal variability, Poisson reduced-rank regression, which models low-dimensional shared structure between neurons. We identified a reliable low-dimensional subspace of activity locally shared among V1 neurons. This subspace represented a large fraction of trial-to-trial spike count variability, and in particular, explained essentially all super-Poisson variability at the population level. Globally distributed patterns of cortical activity, recorded simultaneously from either widefield calcium imaging or with large-scale electrophysiology in frontal cortex, accounted for more than half of this locally explainable variability, significantly more than that attributable to behavior. Optogenetic perturbations suggested a possible causal pathway for these shared fluctuations from the anterior cingulate area. Our results demonstrate that V1 neurons have responses that can be understood as modulated Poisson processes with at least half of the modulations shared beyond V1.

## Results

To determine the magnitude and origin of explainable variability in primary visual cortex (V1), we recorded with a Neuropixels 2.0 probe in awake mice (n=6 sessions, n=6 mice, n=1571 total neurons, 262 ± 34 neurons per session, Figure 1A, B). Subjects passively viewed natural scenes while their face movements were monitored with video cameras (Figure 1C, D). The presentation of images reliably drove responses in V1 neurons. Individual neurons showed tuning to different images (Figure 1E) and trial-to-trial variability, measured by Fano factor (trial-aligned spike count variance to mean ratio), in their responses to identical images (Figure 1F).

**Figure 1:**
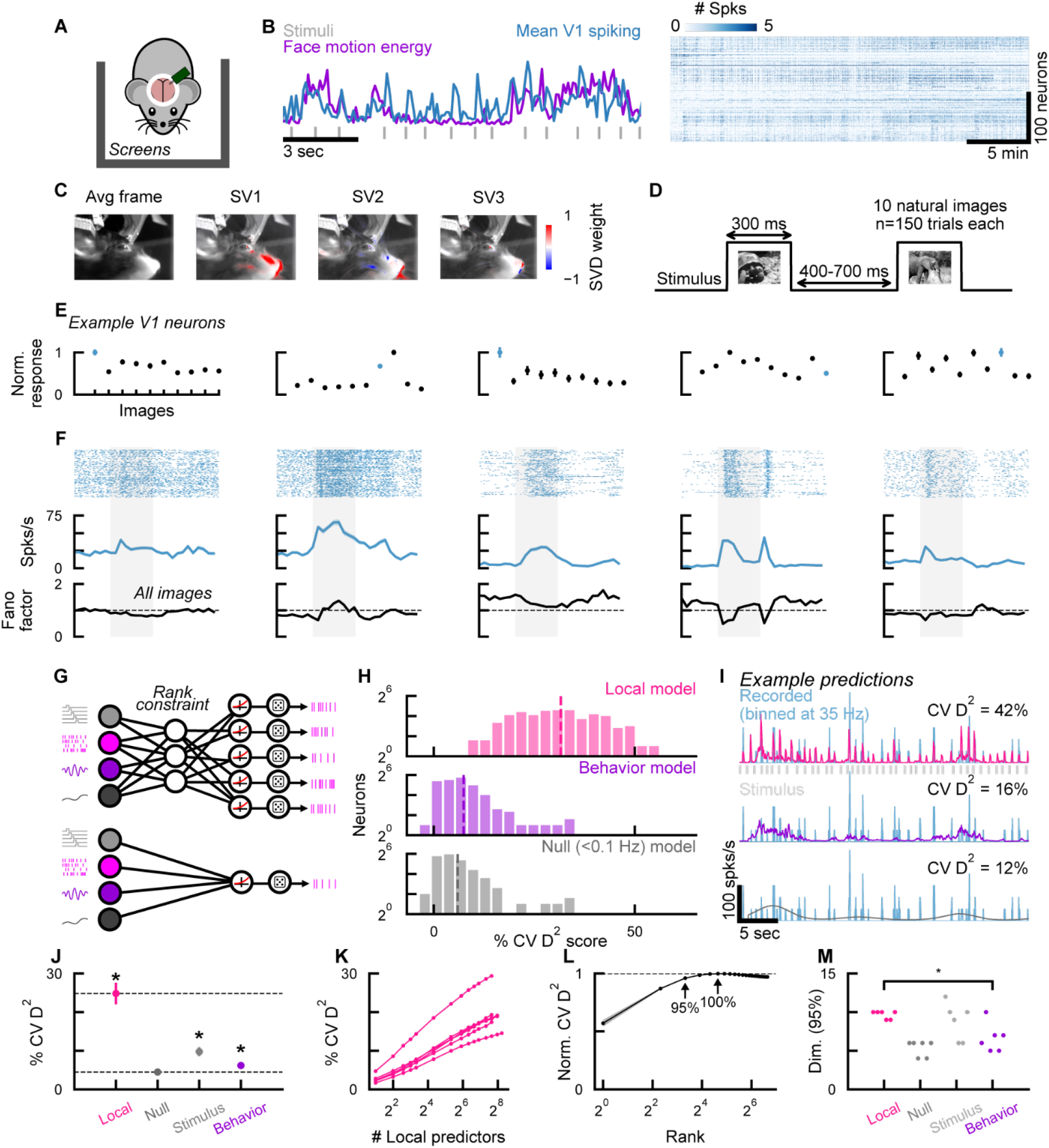
Shared local activity patterns in primary visual cortex explain more spike count variability than behavioral or sensory factors. **A.** Cartoon of Neuropixels 2.0 experimental setup. **B.** Left, example time series of simultaneously collected data. Right, spike matrix from one session binned at 35 Hz (∼28.6 ms), n=220 individual V1 units passing quality control. **C.** Mean frame from a video recording of the subject’s face, and spatial masks corresponding to the top 3 SVD components of the movie, emphasizing parts of the subject’s snout, whisker pad, and eyes. **D.** Image presentation timing. Stimulus identities were randomly interleaved. **E.** Tuning across different images for five example V1 neurons (columns) from one session. Blue indicates stimulus identity for plots in F. **F.** Top, spike rasters corresponding to all trials (n=150) of example stimulus (indicated by blue point in E). Gray bar indicates stimulus duration. Middle, peri-stimulus time histogram corresponding to raster. Bottom, Fano factor of neuron across all ten images. **G.** Schematic of Poisson regression model with a rank constraint to predict multiple targets (top) or without a rank constraint (as a Poisson-GLM) to predict a single target (bottom). Poisson reduced-rank regression nodes from left to right: predictors (stimulus in light gray, spike trains in pink, behavior in purple, null in dark gray), rank constraint, nonnegative nonlinearity function (softplus), Poisson-random noise. **H.** Distribution of prediction scores (cross-validated Poisson deviance explained, or D2) across all neurons of an example session using three different models (Local, Behavior, or Null). Vertical dashed lines indicate means. **I.** Example single-neuron predictions from models in H, overlaid on recorded binned spikes (blue). **J.** Mean D2 scores across n=6 sessions and n=1571 neurons. Asterisks indicate session mean significance relative to Null model (paired t-test across sessions). **K.** Local model performance as a function of the number of peer neurons used as predictors, in a single-target prediction model. Lines indicate sessions. **L.** Local model performance for a multi-target prediction model (averaged across sessions, and normalized to maximum performance for each session) as a function of rank constraint. Shaded error bars indicate S.E.M. across sessions. The rank for 95% performance is 10, and the rank for maximum performance is 25. **M.** Dimensionality, quantified as the rank needed for 95% performance, across model predictors. Asterisk indicates significant difference in dimensionality between models (paired t-test across sessions).

To capture the influence of factors that may explain trial-to-trial variability in V1 spike counts, we developed a Poisson reduced-rank regression (“p-RRR”) model (see Methods). The p-RRR model is a hybrid of regression-based and latent variable models, with a nonlinearity and Poisson “output” step to model observed spike trains. It models a low-dimensional shared latent structure between predictors and targets through its rank constraint, guided by the observation that neural populations often have low-dimensional shared structure (Cunningham and Yu, 2014; Gao and Ganguli, 2015). A rank constraint on stimulus and task variables has been shown to be useful in a previous model for neuron spiking response (Latimer and Freedman, 2023), and p-RRR extends it to model correlations between simultaneously recorded neurons. The use of a Poisson output nonlinearity describes the distributions of spike counts more accurately than a linear model (Semedo et al., 2019), and successfully captured ground-truth latent factors in simulated data (Supp Figure 1A-D). The p-RRR model performed better than the linear model on our dataset (Supp Figure 1E). When used to predict a one-dimensional target, the p-RRR model behaves as a Poisson generalized linear model (GLM) (Paninski, 2004; Truccolo et al., 2004) (Figure 1G). When non-stimulus factors are included alongside stimulus information as predictors, p-RRR models the unobserved source of trial-to-trial variability as a function of these factors and serves as a flexible generalization of the modulated Poisson model (Goris et al., 2014). Specifically, in prior work, shared super-Poisson variability (i.e., variability in excess of that expected from a Poisson process) was modeled as a time-varying quantity of unknown origin, and this model enables identification of the correlates of that shared variability. Besides neural activity, the p-RRR model may also be applicable to other count-based data; a variant of it has been used for analyzing sequencing data (Fitzgerald et al., 2022).

A “peer prediction” of V1 neurons captured a major fraction of the V1 variability via locally shared factors in the neural population. We first used the p-RRR as a single-target GLM, predicting each V1 neuron from the spiking activity of other single neurons in the simultaneously recorded population. This “Local” model explained 24.8 ± 2.8% (mean ± SEM) of the total variance in held-out test data (Figure 1H-J). We used a D^2^ score (Tweedie deviance, similar to R^2^ but for Poisson-distributed model outputs) on held-out data to score each model. The optimal model performance was similar whether we used this metric or R^2^ (Supp Figure 1E), and whether we structured the model to predict single or multiple targets (Supp Figure 2A, B). As a baseline against which to compare model performance, we constructed a “Null” model whose predictors were the slow (<0.1 Hz) components of the population activity (see Methods). The Null model captures slow processes that can result in nonsense correlations (Harris, 2021) and are not the focus of this study, such as representational drift (Deitch et al., 2021), “infraslow” population activity (Okun et al., 2019), or artifactual relative motion between electrodes and the tissue. The Local model predicted significantly more variance than the Null model (p=0.0004, paired t-test across session mean D^2^). Previous work reported that a significant fraction of V1 neural variability is explained by overt movements observable in facial videos (Musall et al., 2019; Stringer et al., 2019b). These factors also explain some V1 spike count variability in our dataset (“Behavior” model), but leave a major fraction unexplained (Figure 1J). The Behavior model explained significantly less spike count variability than the Local model (p=0.0008) but significantly more than the Null model (p=0.002, Supp Figure 3A).

Spike count variability was accounted for by a low-dimensional set of shared local factors. While performance did not saturate with increasing numbers of neurons used to make predictions (Figure 1K), this lack of saturation can arise due to spiking noise even in low-dimensional simulated populations (Supp Figure 2C, D). In other words, improved performance with more neurons can arise from a better estimation of low-dimensional shared variables, and therefore does not necessarily indicate a population dimensionality higher than the number of recorded neurons. We more directly estimated the dimensionality via the reduced rank structure of our multi-target model, determining the rank needed to reach at least 95% of maximum performance (Figure 1L). Measured in this way, the dimensionality of spike count variability of local V1 activity was 9.8 ± 0.18 (mean ± S.E.M. across sessions), significantly greater than the dimensionality explainable by correlations with behavior (6.6 ± 0.7; Figure 1M; p=0.009, paired t-test) indicating that the full space of V1 activity explores dimensions not associated with behavioral factors.

Accounting for locally-shared variability revealed that, on average, V1 neurons fired with variability close to a Poisson process. We quantified the total trial-to-trial spiking variability with Fano factor (Figure 2A). The median Fano factor of V1 neurons across sessions was 1.17 ± 0.10 at baseline (median ± median absolute deviation across n=6 sessions), decreasing to 1.12 ± 0.10 immediately following stimulus onset (Figure 2B, paired t-test p=0.01), a decrease that was consistent with prior observations in other species (Churchland et al., 2010). The Local model captured much of this trial-to-trial variability, correctly predicting differences in amplitude of responses around the time of stimulus presentations (Figure 2C, E).

**Figure 2:**
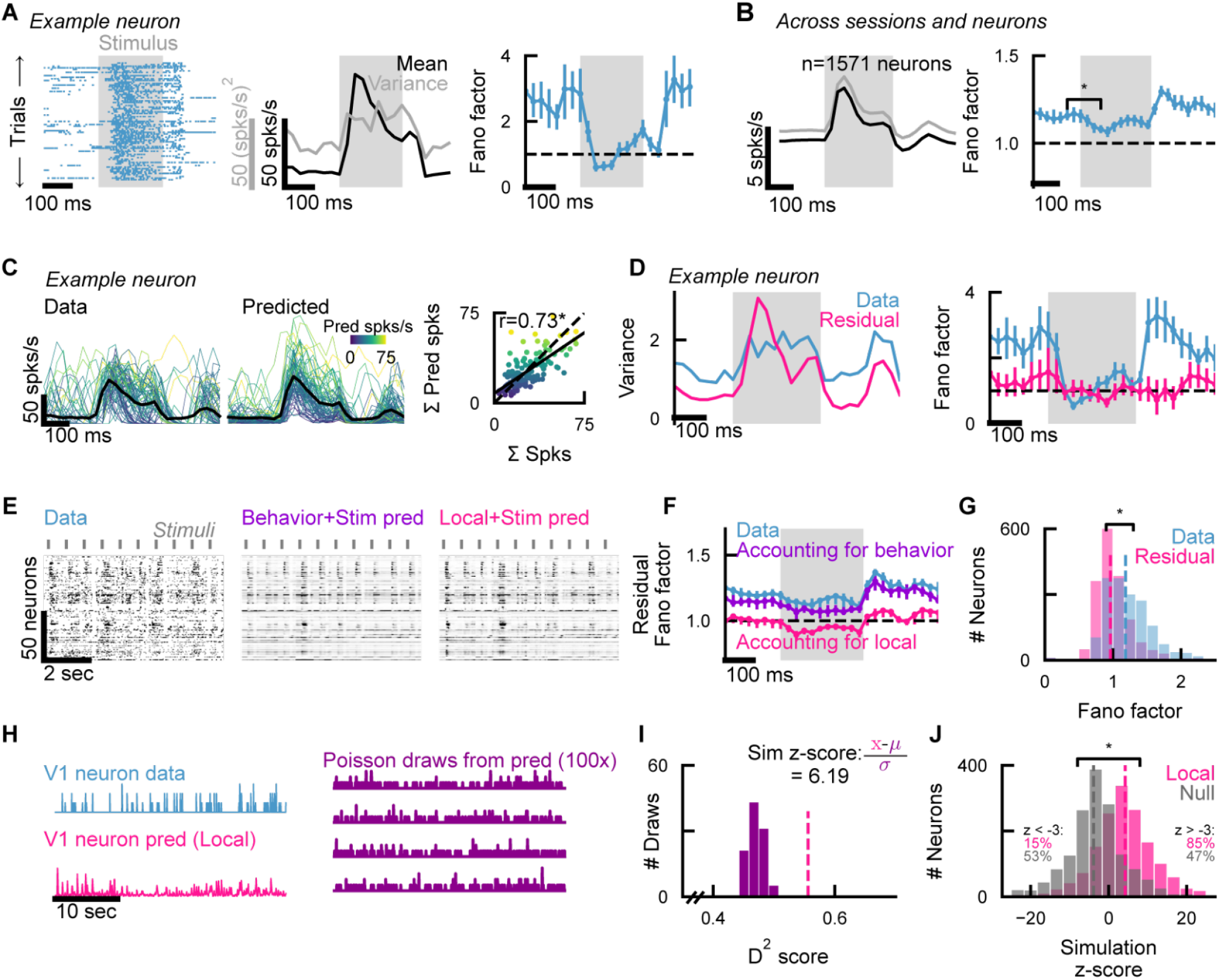
Shared factors within local populations explain super-Poisson components of trial-to-trial variability. **A.** Example Fano factor calculation for a single neuron. Left, spike rasters for all trials of an identical stimulus. Middle, mean (black) and variance (gray) of spike counts calculated across trials in 35 Hz (∼28.6 ms) bins. Right, single neuron Fano factor, i.e. the spike count variance to mean ratio. Error bars indicate 95% confidence intervals from bootstrapping within trials. **B.** Left, mean (black) and variance (gray) across ten image conditions for n=1571 neurons from n=6 sessions. Right, median Fano factor across sessions. Error bars indicate 95% confidence interval obtained from bootstrapping within sessions. **C.** Left, example single-trial responses of example V1 neuron from one stimulus. Right, predicted versus recorded spike counts for individual trials (Pearson correlation coefficient, p<1e-8). Colors indicate the sum of predicted spikes on each trial. Black line indicates mean across trials. **D.** Left, recorded spike count variance and residual (i.e. recorded minus predicted) spike count variance for neuron in C. Right, recorded and residual Fano factor (variances, plotted left, divided by recorded means). **E.** Example population activity, recorded and predicted, for one session. **F.** Original and residual Fano factors across sessions. Error bars indicate 95% confidence intervals from bootstrapping within sessions. **G.** Distribution of recorded (1.18 ± 0.22, median across all timepoints and neurons ± median absolute deviance) and residual Fano factor (0.96 ± 0.14; p<1e-8, Wilcoxon signed-rank test). **H.** Schematic of Poisson simulation test. Poisson-random time series were taken from the model prediction (i.e., the predicted value at each time point was used as the mean parameter for a draw from a Poisson distribution). **I.** Simulation and scoring for an example neuron. Purple distribution represents D2 scores between the prediction and Poisson draws (pink and purple in H). Pink line represents the D2 score between the data and prediction. Simulation z-score for example neuron is 6.19. **J.** Distribution of simulation z-scores computed from the Local model prediction and the Null model prediction. (p<1e-8, paired Kolmogorov-Smirnov test).

To assess how close the remaining unexplained variability was to that expected from a Poisson process, we computed a “residual” Fano factor (variance of the difference between observed and predicted single-trial spike count data, divided by the observed trial-average spike count, Figure 2D, see Methods). This “floor” of unexplainable variability has a Fano factor of 0.99 ± 0.03 (mean ± 95% confidence interval obtained from bootstrapping, Figure 2F), significantly lower than the raw Fano factor of 1.15 ± 0.04 (Figure 2F, paired t-test p<0.0001). We refer to this as the “floor” since, given that most reliable V1 variance is captured by a small number of dimensions, most explainable variability should be shared across neurons and thus captured by this model. As with the total predicted variance (CV D^2^ in Figure 1L), the residual Fano factor did not saturate as a function of the number of local peer predictors (Supp Figure 4B). We noted that individual neurons were heterogeneous in their spiking statistics, with some neurons far above or below Poisson expectation (Figure 2G), despite the overall population average being close to the variability expected from a Poisson process. This individual neuron spiking variability was uncorrelated with spike rate, spike waveform width, and unit depth (Supp Figure 5A).

Despite its widespread use, the Fano factor also has challenges as a reliable indicator of Poisson-like spiking variability (Charles et al., 2018; Churchland et al., 2011), including its limitation to trial-aligned activity. We therefore developed a complementary, alternative analysis using simulated Poisson counts. This method directly examines the relationship between the residual variability observed for each individual neuron and the variability expected from a Poisson process. For each neuron, we generated 100 Poisson-random samples from the Local or Null model predictions (Figure 2H). We then scored these samples against the prediction, creating a distribution of D^2^ scores expected under conditions of perfectly accurate mean rate predictions (by construction) combined with exactly Poisson spike generation (Figure 2I). Finally, we computed a z-score for each neuron’s actual D^2^ score relative to this distribution. A z-score of zero indicates that the model predicted the same amount of variability that would be expected if the neuron were a Poisson spike generator with the predicted mean rate. Positive z-scores indicate more variability predicted, thus implying that the neuron fires more regularly than Poisson given the model prediction, and vice-versa for negative z-scores. The results from this method were reliable and robust to cross-validation (Supp Figure 5B). As expected, when the Null model was used for prediction, most neurons had significantly (z<-3, 99.7% confidence interval) negative z-scores indicating their super-Poisson variability (55% of neurons with z<-3). However, when the Local model was used for prediction, a substantial fraction of the population had responses that were at or below Poisson variability (84% of neurons with z>-3). These simulations demonstrate that most neurons fired similarly to a Poisson process or with somewhat more regular firing, but only after accounting for locally-shared variability (Figure 2J).

In principle, the locally-shared activity patterns that we identified could be partially or completely isolated in V1, or may be shared with fluctuations in other parts of cortex. In order to identify a potential source for globally-shared activity, we mapped causal cortical inputs to V1 using simultaneous widefield imaging and scanning optogenetics (Matveev et al., 2024). The anterior cingulate area in mice projects directly to V1 (Zhang et al., 2014), and indeed we observe that suppressing activity there directly resulted in suppression of V1 responses (Figure 3A-C). Therefore, we hypothesized that anterior cingulate area (ACA) might be a conduit of globally shared activity underlying trial-to-trial variability V1. While recording in V1, we measured nonlocal (“Global”) cortical activity using either a second Neuropixels 2.0 probe in anterior cingulate area (ACA; n=3 sessions; Figure 3D-E), or widefield calcium imaging of the cortical surface (n=2 sessions; Figure 3F-G), or both methods simultaneously (n=1 session). We fit the p-RRR model to predict V1 using this Global activity, face movements, stimulus kernels, and the null term (Figure 1H). This “Full” model achieved just over half of the predictive power of the Local model (12.4 ± 1.0% for Full vs 24.8 ± 2.8% for Local, Figure 3I). Adding Global predictors to the Local model did not improve performance (Supp Figure 4A), consistent with the assumption that all V1 signals arising from external sources are also shared locally among neurons within the V1 population.

**Figure 3:**
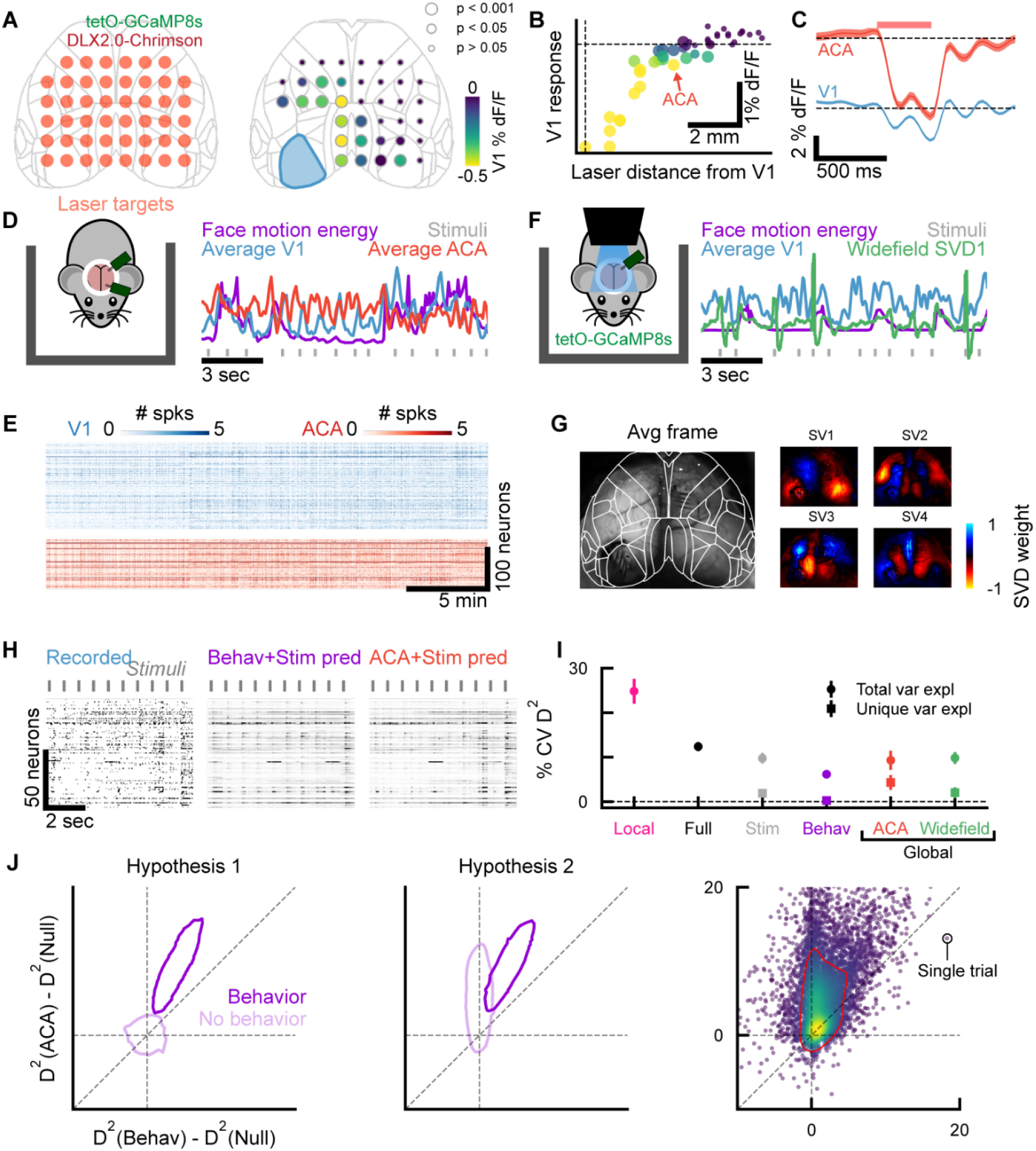
Globally-shared activity patterns reflect nearly all behavioral components of local spike count variability. **A.** Left, schematic of causal pathway mapping via cortex-wide optogenetics (via injected DLX2.0-Chrimson) and calcium imaging (via transgenic GCaMP8s). Right, map of causal pathways to V1. Circle colors indicate mean response in V1 following inhibition at corresponding laser target (n=620 trials over n=8 sessions). Areas adjacent to V1 are removed for visual clarity (but see **B**). **B.** Magnitude of downstream inhibition in V1 as a function of optogenetic target distance. Dashed lines indicate zero. Size corresponds to significance test as in **A.** **C.** Timeseries of V1 and ACA calcium response during ACA-targeted inhibition (±0.75 ML, +0.5 AP; n=400 trials from example session). **D.** Left, cartoon of dual-probe Neuropixels 2.0 recording. Right, example time series of simultaneously collected data. **E.** Heatmap of neuronal activity (binned at 35 Hz, ∼28.6 ms) simultaneously recorded from two populations. **F.** Left, cartoon of widefield calcium imaging paired with Neuropixels 2.0 recording. Right, example time series of simultaneously collected data. **G.** Example mean frame from widefield calcium imaging. Spatial masks correspond to the top four components of the widefield movie. **H.** Example population predictions with Behavioral and Global predictors. **I.** Cross-validated Poisson deviance explained for different predictors (circles). Squares indicate unique variance explained (relative to “Full” model). **J.** Left and middle, schematic of two hypotheses for single-trial results. In Hypothesis 1, global signals only carry behavioral information. In Hypothesis 2, global signals also carry non-behavioral information. Right, prediction score from Nonlocal model versus prediction score from Behavior model on individual trials. Points are colored by local density. Red line indicates density contour.

A “Global” model, which included nonlocal activity and the null term but not behavior or stimulus kernels as predictors, was significantly better than the Behavior model in predicting V1 spike counts (6.2 ± 0.8% for behavior, 9.3 ± 2.1% for ACA (paired t-test p<0.0001), 9.8 ± 1.3% for widefield (paired t-test p<0.0001), Figure 3I). Moreover, when Global predictors were included in the model, Behavior predictors did not add explanatory power. We computed the unique contribution of each predictor by removing it from the Full model and quantifying the reduction in performance. The unique contribution of behavior was nearly zero (0.3 ± 0.3%).

The lack of unique explained variability for the Behavior model relative to the Global model was unlikely due to undersampling of the behavioral space. In prior work, it had been observed that cortical population activity predictions were minimally improved with additional videos of the mouse beyond the face, due to the high correlation of bodily with facial movements (Manley et al., 2024). In principle, both ACA and V1 could receive copies of the same behavioral signal that give rise to the ability of ACA to predict V1. Our optogenetic experiments demonstrate that arbitrary non-behavioral signals injected into ACA do transmit to V1, confirming the possibility that activity patterns in ACA unrelated to behavior may directly contribute to variability in V1. We recorded both the subject’s body and face in a subset of sessions, and also found that body movements were highly correlated with face movements (Supp Figure 3B, C). To test this another way, we considered that if Global model predictions of V1 truly only reflected behavioral signatures, then this information should not be predictive of V1 activity on trials when mice are sitting very still with no overt behaviors (Figure 3J, Hypothesis 1). Under this hypothesis, higher predictability with the Global model reflects better predictions than the Behavior model on the same trials, potentially arising from a higher-resolution representation of the same behavioral information. If instead the Global activity carried both behavioral and other, non-behavioral predictive signals, then the trials without behavior would still be predictable from the Global model (Figure 3J, Hypothesis 2). We found that trials in which behavior held no predictive power were nevertheless predictable from global activity, consistent with Hypothesis 2. The superior predictability of V1 activity by the Global model relative to the Behavior model was consistent across synchronized and desynchronized cortical states (Supp Figure 5C, D). These results demonstrate that the predictions of V1 activity by global signals include fundamentally distinct factors rather than being a more detailed representation of behavior alone.

A geometric analysis of predicted V1 manifolds reinforced the conclusion from the previous D^2^ comparison. In this analysis, we asked whether the Global model explains shared V1 variability along unique dimensions in V1 population activity relative to the Behavior model. Following previous studies (Cunningham and Yu, 2014; Gallego et al., 2018, 2017; Sadtler et al., 2014), we took V1 neural space as a high-dimensional space where each axis represents the activity of one neuron. For all models, both the optimal model rank and prediction dimensionality were significantly lower than the number of simultaneously recorded neurons (Figure 4A-C), suggesting that the predictable activity mainly evolves on low-dimensional manifolds of the neural space. To identify the manifold for each model, we applied principal component analysis (PCA) to the model prediction, and defined its manifold as the *r*-dimensional subspace spanned by the top *r* PCs capturing over 99% variance (Gallego et al., 2018). To quantify the alignment between the Global and Behavior manifolds, for example, we would compute the percentage of variance of the Global model prediction that was captured within the Behavior manifold. Larger values would indicate that more variance is captured by shared neural modes between the two manifolds, and thus a higher degree of alignment.

**Figure 4:**
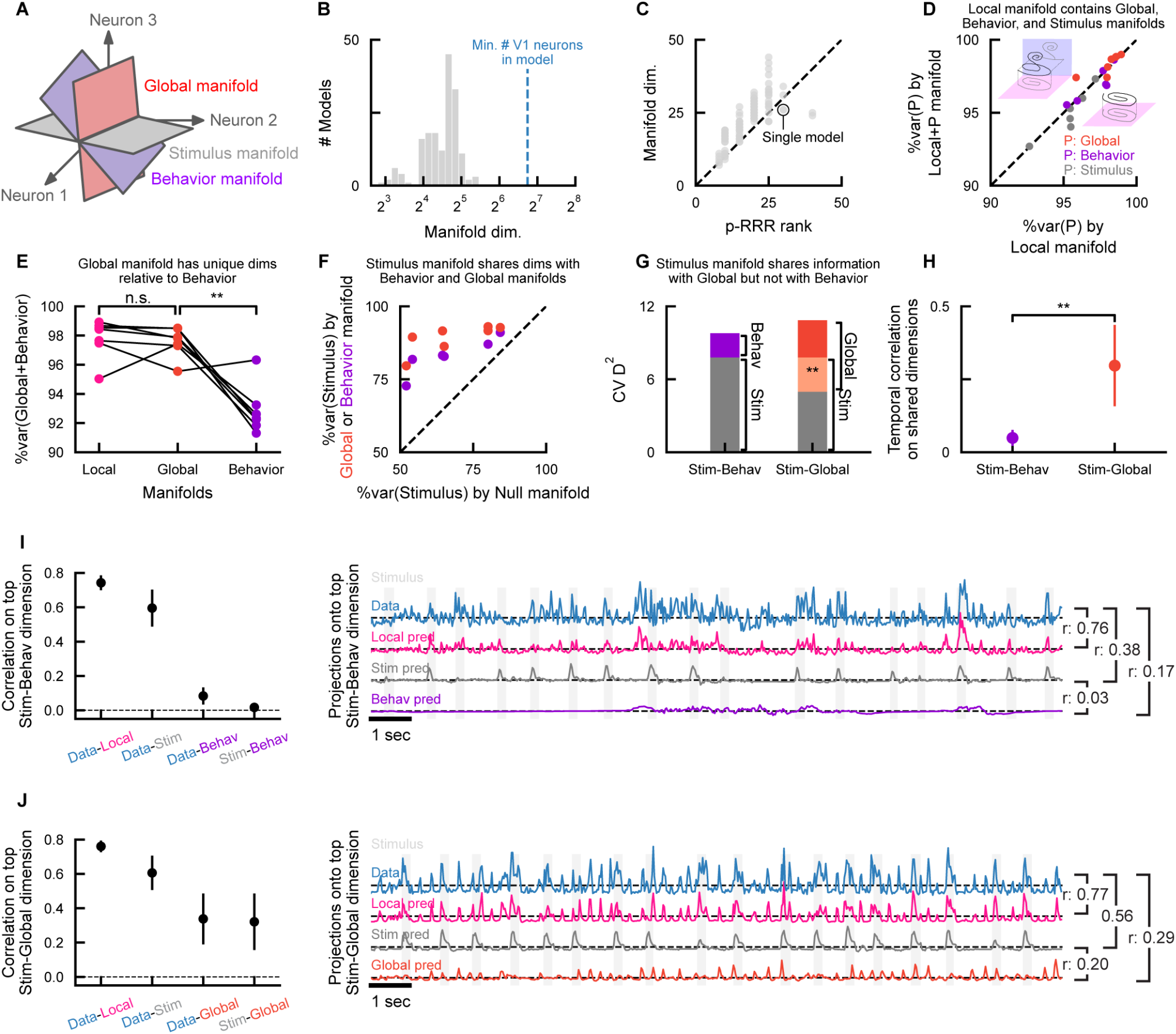
The V1-Behavior manifold is a subspace of the Global manifold, and shares dimensions but not temporal information with the Stimulus manifold. **A.** Schematic of predicted low-dimensional manifolds in the V1 neural space. **B.** Dimensionality of predicted manifolds, pooled over all sessions and all models (Null, Stimulus, Behavior, Global, Local, and their combinations; n=172). **C.** Dimensionality of p-RRR predicted manifolds versus p-RRR model ranks, pooled over all sessions and all models. Predicted manifolds are of slightly higher dimensionality than p-RRR rank due to the rectifying nonlinearity in p-RRR. **D.** Nonlocally (using Stimulus, Behavior, or Global, as indicated by colors) predicted variance accounted for by manifolds predicted by combined Local and nonlocal predictors (y-axis), versus by local peer predictors alone (x-axis). Insets: cartoons illustrating nonlocally predicted variance completely (right) or partially (left) captured by Local manifold. Black curve: nonlocally predicted activity. Pink plane: Local manifold. Purple plane: nonlocally predicted manifold. **E.** Variance predicted by combined behavior and global neural activity accounted for by Local, Global, or Behavior manifolds. **F.** Stimulus predicted variance accounted for by Behavior or Global manifolds (y axis), versus by Null manifold (x axis). Recall that the Null term is present in all Stimulus, Behavior, and Global models. **G.** Unique and shared *D*^2^ between Stimulus and Behavior models, and Stimulus and Global models. The contribution of the null term is subtracted when computing shared *D*^2^: *D*^2^(Stim+Behav)=*D*^2^(Null+Stim+Behav)-*D*^2^(Null), *D*^2^(Stim)=*D*^2^(Null+Stim)-*D*^2^(Null), *D*^2^(Behav)=*D*^2^(Null+Behav)-*D*^2^(Null), and shared *D*^2^(Stim, Behav)=*D*^2^(Stim)+*D*^2^(Behav)-*D*^2^(Stim+Behav). *D*^2^(Stim+Global), *D*^2^(Global), and *D*^2^(Stim, Global) are computed similarly. **H.** Correlation between projected activities along shared dimensions. To find shared dimensions beyond those driven by the Null term, the Null manifold is first projected out from the Behavior manifold via an orthogonal projection. The shared dimensions are then identified as dimensions in the Behavior manifold that successively maximize stimulus-related variance (Methods). Both Stimulus and Behavior predictions are then projected onto these dimensions to obtain projected activities. Correlation is computed between projected activities onto each shared dimension, and then averaged across all shared dimensions, weighted by stimulus-related variance captured on individual dimensions. Shared dimensions with Global manifolds are found similarly. Correlation on stimulus and behavior shared dimensions is significantly smaller (n=7, p=0.0078, one-tailed Wilcoxon signed-rank test). **I.** Correlation between projected activities along the top stimulus-behavior shared dimension, which is the dimension within behavior predicted manifold that captures the most stimulus-related variance. Recorded V1 activity (Data), Local model prediction, Stimulus model prediction, and Behavior model prediction are then projected onto the top dimension, and correlations are computed between projected activities. Projected activities in one example session (as indicated by cyan markers) are visualized. Shaded regions indicate stimulus presentations. **J.** Similar to I, with Behavior model substituted by Global model.

Following our strategy above, we first determined the relationships with the manifold of local predictions (Local manifold). Consistent with results from the unique variance analysis (Supp Figure 4A), we found that the Global, Behavior, and Stimulus manifolds were subspaces of the Local manifold. Combined Local+Global, Local+Behavior, and Local+Stimulus manifolds explained similar amounts of variance compared to the Local manifold alone (Figure 4D, p>0.1 for all predictors, two-sided paired sample t-test), suggesting as expected that these predictors do not drive unique dimensions outside of the Local manifold.

Next, we showed that the Global manifold contained the Behavior manifold, and extended into unique dimensions. Specifically, we took the prediction made by the combined Global+Behavior model, and computed the percentages of its variance captured by the Global or Behavior manifolds alone. All of the combined model variance was captured within the Global manifold, that is, was indistinguishable from the upper bound established by the Local manifold (p=0.66, two-sided paired sample t-test). By contrast, the portion of combined model variance captured within the Behavior manifold was significantly smaller (p=0.0015, one-tailed paired sample t-test) (Figure 4E). Therefore, the Global model predicts unique dimensions that are not predicted by the Behavior model, but not vice versa. While this may be unsurprising given the unique variance analysis (Figure 3F), it does not follow trivially from it, as it is possible for the two different predictors to encode distinct temporal patterns along shared dimensions. Interestingly, we found this to be the case for modes shared by the Behavior and Stimulus manifolds. While they share dimensions beyond those in the Null manifold (Figure 4F, p=0.0022 for Behavior, p=0.0008 for Global, one-sided paired sample t-test), in terms of *D*^2^ there appears to be minimal overlap between the information they carry (Figure 4G, shared *D*^2^ = 0.000±0.003, p=0.53, one-tailed one sample t-test). Examining their projected activity along the shared dimensions, we found that they largely account for distinct aspects of V1 fluctuations, and show little temporal correlation (Figure 4H, I). In contrast, when comparing Stimulus and Global models, we observed correlated activity along shared neural modes (Figure 4H, J), as expected given by their significant overlap in *D*^*2*^ (Figure 4G, shared *D*^2^ = 0.028±0.014, p=0.0014, one-tailed one sample t-test). Together with results from the explained variance comparison (Figure 3), these findings suggested that most of the behavior-related V1 variability can also be found among shared fluctuations coordinated across the brain.

## Discussion

Using a novel approach for partitioning spike count variability that allows identification of sources, we discovered that a low-dimensional subspace of activity within V1 accounts for super-Poisson components of trial-to-trial variability. Over half of this explainable variability was attributable to globally distributed cortical activity patterns. The behavioral factors were contained in the globally-predicted subspace, which could predict V1 activity even on trials without overt behavior. Whereas local recurrent connectivity has been assumed to underlie dynamic computations at the level of individual regions (Brunel and Wang, 2001; Rubin et al., 2015), our result that a majority of fluctuations are shared suggests instead that many computations may be fundamentally distributed across brain regions.

Our model makes several key assumptions. First, the model assumes that neuron spike counts are described by a linear combination of predictive factors with a nonlinearity after summation. While such models have provided good fits to spike count data in a variety of contexts (Musall et al., 2019; Park et al., 2014; Pillow et al., 2008), we have not explored the extent to which model fits could be improved with, for example, nonlinear interactions between predictors. Second, we assume that predictive relationships are static over time.

However, V1 stimulus representations may drift on slow timescales (Deitch et al., 2021) and cortical dynamics may be better accounted for by switching dynamical systems (Akella et al., 2025; Linderman et al., 2016). In the future, models that allow for non-stationarity in predictive relationships may improve fits. Due to these two assumptions, our model represents only a lower bound of the possible predictive performance. Nevertheless, our Local model already captures essentially all super-Poisson variability. Considering the Global model, if a version were devised with substantially superior performance, either due to improved model design or extended measurements of global neural activity, this would only strengthen our conclusion that a majority of spike count variability is shared globally. Finally, we assumed a Poisson noise model, which is a common and simple assumption for spike count distributions (Goris et al., 2014; Pandarinath et al., 2018; Yu et al., 2009). This noise model does not prevent us from discovering responses that have less than Poisson variability, and indeed we did find some such neurons (Figure 2G, J).

The population average of unexplainable variance was approximately equal to the mean activity of recorded neurons (Figure 2F), consistent with irreducible Poisson variability combined with time-varying underlying firing rates (Shadlen and Newsome, 1998). This conclusion is consistent with previous observations that visual cortical counts are well described by a modulated Poisson model (Goris et al., 2014). Notably, in that work, rate inhomogeneity of the Poisson process was captured with a multiplicative gain factor, whereas here we capture rate fluctuations using a nonlinear transformation of additive contributions from identified sources. These different modeling choices imply different underlying mechanisms by which shared factors influence neuronal activity, but since the quality of fit appears to be similar with each, more mechanistic methods may be required to disambiguate.

Though there is ample evidence for contributions to neural activity that do not conform to the rate-based coding scheme, such as oscillatory phase locking (Buzsáki and Draguhn, 2004; Fries et al., 2001) and spike latency coding (Kayser et al., 2009; VanRullen et al., 2005), for the timescale and context of our measurements we find close agreement with the predictions of rate coding. Nevertheless, a subtlety of our results represents a key deviation from this model: while the population average of unexplainable variance was close to Poisson, this property was highly heterogeneous across neurons, with some neurons exhibiting reliably less unexplainable variance and others reliably more (Figure 2G, J). One possibility to explain this outcome is that different cell types have different levels of explainable spike count variability. We separately analyzed neurons with broad and narrow waveform durations, and did not observe significant differences in this property between the two classes (Supp Figure 5A). A finer division by cell type identity in the future, using opto-tagging, could resolve this question further. A more intriguing possibility is that Poisson variability is enforced by mechanisms at the population level, with individual neurons deviating from this pattern to reflect specific roles in the network.

We found that the pre-stimulus trial-to-trial variability of mouse cortex is similar to that of primate cortex (Fano factor = 1.0-1.5), and shares a similar “quenching” property after stimulus onset (Churchland et al., 2010). This reduction in variability can theoretically arise from network effects such as an input-stabilized cortical network (Hennequin et al., 2018), a chaos-suppression network effect (Rajan et al., 2010), or a clustered multi-attractor circuit (Litwin-Kumar and Doiron, 2012). Here, variability quenching in V1 persisted even after accounting for locally shared activity patterns (Figure 2F). These results suggest that quenching in V1 may occur primarily through cell autonomous mechanisms such as shunting inhibition (Borg-Graham et al., 1998), rather than through correlations among neurons in the local network. Alternatively, it may be a high-dimensional network effect that our low-rank model does not capture.

Prior work in primates has shown that trial-to-trial spiking variability is low when behavior is controlled (Gur et al., 1997), suggesting a similar contribution of behavior-related variability as what we observed in mice. However, behavioral correlations with cortical activity may be weaker in primates compared to mice, and more of these fluctuations may be related to aspects of internal state such as arousal (Liska et al., 2024; Talluri et al., 2023). This difference suggests species-specific computational functions for these correlated activity patterns, potentially reflecting different behavioral priorities for vision across species (Kang et al., 2025). At least some V1 variability in primates is correlated with other cortical regions such as V2 (Semedo et al., 2019) and V4 (Semedo et al., 2022). Whether a majority of V1 variability in primates is predominantly confined locally or shared globally, as in our data from mice, remains an open question.

Here, we demonstrated that locally-shared activity in V1 explained nearly all super-Poisson trial-to-trial variability, but globally-shared activity accounted for the majority of that. This finding suggests that private fluctuations within the V1 recurrent circuit are limited to less than half of the total space of V1 activity. The global cortical measurements in this study captured just a subset of all cortical activity patterns, so future studies that simultaneously record from more regions may reveal that nearly all super-Poisson variability in V1 is shared across the mouse brain. Nevertheless, our results demonstrate that distributed computation dominates private computation in visual cortex.

## Acknowledgements

We thank Ljuvica Kolich and Kimberly Miller for help with mouse husbandry; Greg Horwitz, Nathan Kutz, Fred Rieke, and Brent Doiron for helpful feedback; Joseph Pemberton and Zhiwen Ye for comments on the manuscript. The optogenetic virus was provided by Jonathan Ting and Ximena Opitz-Araya (Allen Institute for Brain Science).

This work was supported by the National Science Foundation (NCS-FO 2024364 to E.S.B. and N.A.S.), the NIH Ruth Kirchenstein award (T32EY007031 and F31EY035880 to A.J.L), the Simons Foundation Shenoy Undergraduate Research Fellowship (to P.M.), the Pew Biomedical Scholars Program (to N.A.S.), and the Klingenstein Fellowship in Neuroscience (to N.A.S.).

## Author Contributions

A.J.L., Z.L., E.S.B., and N.A.S. conceived and designed the study. A.J.L performed electrophysiology experiments. A.J.L, A.E.L., and P.M. performed optogenetic experiments. Z.L. developed the model. A.J.L. and Z.L. performed data analysis. A.J.L., Z.L., and N.A.S. wrote the original draft manuscript. All authors revised the manuscript. N.A.S. and E.S.B. supervised and obtained primary funding for the project.

## Data Availability

Data will be available in NWB format on the DANDI archive prior to publication.

## Materials and Methods

### Animals

Experiments were performed on male and female mice single housed on a 12 hour light/dark cycle. For experiments with widefield imaging, multiple GCaMP-expressing genotypes were used: either TetO-G6 s;Camk2a-tTa (B6;DBA-Tg(tetO-GCaMP6s)2Niell/J or TetO-G8s;Camk2a-tTA (B6;*D*^2^-Tg(tetO-GCaMP8s)1Genie/J crossed with B6.Cg-Tg(Camk2a-tTA)1Mmay/DboJ). Wild-type (C57Bl6/J) mice were used for experiments with only electrophysiology.

### Surgery

An initial surgery was performed under isoflurane (1-4% in O2). Animals were given carprofen (5 mg/kg) subcutaneously in the shoulders and lidocaine (2 mg/kg) subcutaneously in the surgical area above the skull. The skin was cleared to reveal the surface of the dorsal skull. The edges of the incision were glued to the skull with cyanoacrylate (VetBond, World Precision Instruments) to protect the underlying muscle. The skull was treated with 10% citric acid (Dentin Activator Liquid, Parkell) to improve the bond between skull and implant. Then, a 3D-printed recording chamber was secured to the skull with dental cement (Metabond, Parkell). Thin layers of UV-cured optical glue (Norland Optical Adhesive 81, Norland Products) were applied to the exposed surface of the skull inside the chamber. A 3D-printed titanium headpost (ProtoLabs) was then cemented to the posterior end of the recording chamber and cured successively. After surgery, mice were treated with carprofen for two days and given at least one week to recover before experiments. At least two days before experiments, mice were acclimated to handling and head fixation, and lightly water restricted.

Before electrophysiological recordings, another brief (∼ 20 min) surgery was performed under light isoflurane (1-2% in O2) to open craniotomies for probe insertions. Craniotomies were approximately 2 mm in diameter and performed with a handheld dental drill. Exposed dura was sealed with a silicone gel (Dow Corning 3-4680 Sil Gel), and the chamber covered with a 3D-printed cap to protect the craniotomies. Mice were allowed to recover for at least two hours before experiments.

For optogenetic experiments (Matveev et al., 2024), we retro-orbitally injected tetO-GCaMP8s mice aged P28-42 with the DLX2.0-ChrimsonR-tdTomato opsin packaged in a PHP.eB serotype AAV. Injections were 1.6e12 GC diluted in 100 uL PBS.

### Widefield calcium imaging

Images were collected with a CMOS camera with 17.3 um/pixel resolution (Basler acA2440-75m) and 560 × 560 pixel frame size. The camera was fitted with a 0.63x objective lens (Leica Planapo 0.63x). The true frame rate is 70 Hz, but the effective rate is 35 Hz due to alternating 405 nm and 470 nm excitation for hemodynamic correction. The LED onset and offset were modified to be a 4-ms sine-wave ramp to reduce light artifacts for Neuropixels recordings. To avoid light contamination from the stimulus screens, a 3D-printed black cone was affixed to the top of the recording chamber with Blu-Tack. The top of the cone was attached to a piece of light-blocking fabric (ThorLabs) that rested on the top edge of the screens. Light path details and image preprocessing are as described in Matveev et al. (2024).

### Optogenetics

For cortex-wide mapping experiments (Figure 3A, B), laser targets were 46 coordinates spaced 1 mm apart. Laser duration was 25 ms and amplitude was 0.75 mW. Laser targets were randomly interleaved, with 250-350 ms inter-trial intervals. For ACA-targeted optogenetic experiments (Figure 3C), the laser coordinate was ±0.75 mm ML, +0.5 mm AP relative to bregma. Laser duration was 100 ms and amplitude was 1 mW.

### Electrophysiology

Recordings were made with Neuropixels 2.0 electrode arrays, which have 384 selectable recording sites from 5210 total sites distributed across four 1 cm shanks (Steinmetz et al., 2021). Shanks are 250 um apart and each have two vertical columns of sites. Recording configuration was across all four shanks, and spanning either a depth of 700 um or 1400 um. Probes were mounted to a dovetail rail and attached to a steel rod for micromanipulation (Sensapex). Probe trajectories were planned with Pinpoint (Birman et al., 2023). For later track localization, probes were coated in DiI (ThermoFisher Vybrant V22888) before insertion. In some experiments, probes were inserted through the silicone gel, a saline bath was used in others. Probes were lowered to their final positions at approximately 5 um/s. Recordings were made with 30 kHz sampling. In some cases, additional recordings were made at the same locations on subsequent days. Spike sorting was performed with Kilosort 2.5 using default parameters (Steinmetz et al., 2021). After automatic sorting, units were checked manually in Phy (https://github.com/cortex-lab/phy) for any noise artifacts. We then applied a refractory period correction and a noise amplitude correction. These metrics passed 29.2% of units relative to initial yield.

### Passive stimulus presentation

During each experimental session, mice were presented with visual stimuli on three 60 Hz refresh rate screens (Adafruit, LG LP097QX1). Screens were placed 10 cm in front of the mouse, to its left, and to its right. Stimuli were ten natural scenes from the Allen Brain Observatory Visual Coding dataset (de Vries et al., 2020). Images were shown for 300 ms, and in some sessions, for 230 ms. Inter-trial intervals were exponentially random, with a mean of 700 ms, minimum of 400 ms, and maximum of 3000 ms. To keep mice awake and alert throughout visual stimulus presentations, they were randomly rewarded (on average, once every five seconds) with drops of 10% sucrose delivered through a small metal tube positioned near the mouth.

### Behavioral monitoring

Mice were able to turn a wheel freely during recording experiments. The wheel was attached to a rotary encoder to record movements. A camera and 16 mm focal length lens (ThorLabs MVL16M23) with IR filter was used to record the subject’s face and forelimb movements during experiments. The camera acquired at a 70 Hz frame rate (∼14.3 ms), using the same hardware trigger as for the widefield imaging camera.

### Poisson reduced-rank regression

#### Model fitting and relationship to Poisson GLM

Let 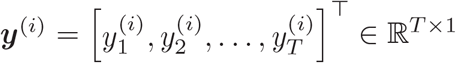 denote the binned spike train of neuron *i*, and ***X*** = [***x***_1_, ***x***_2_,…, ***x***_*T*_]^*Τ*^ ∈ ℝ^*T×M*^ denote the *M* -dimensional measurements (e.g. sensory inputs, behavioral or neural features) recorded at time bins 1, 2,…,*T*. p-RRR assumes that 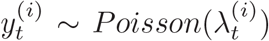, where 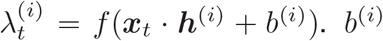is an intercept term accounting for the average activity, and *f* is a nonnegative nonlinearity, which is taken to be the softplus function (Keeley et al., 2020). Once learned, the model parameter, ***h***^(*i*)^ ∈ ℝ^*M×*1^ specifies the relationship between the observed spike counts and the other measurements.

To learn ***h***^(*i*)^, *b*^(*i*)^ from data, we can write down the log-likelihood of the spike train

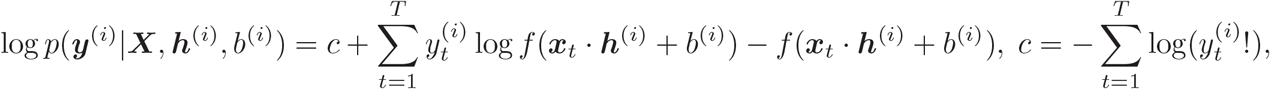

and the optimal ***h***^(*i*)^, *b*^(*i*)^ are those that maximize the likelihood of the observed spike train.

When spikes from multiple neurons are recorded, for example, *i* = 1,…,*N*, the joint log-likelihood for observing all spike trains is

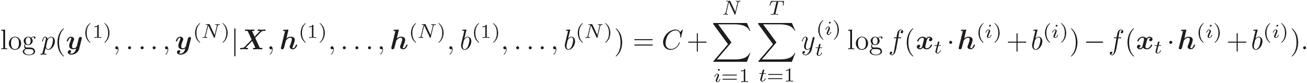

Assuming that 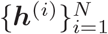are independent of each other, previous approaches maximize the joint log-likelihood by maximizing log *p*(***y***^(*i*)^ | ***X, h***^(*i*)^, *b*^(*i*)^) for each *i* separately. However, given the significance of low-dimensional structure in neural populations, instead of independence, it may be more appropriate to assume ***H*** = ***h***^(1)^, …, ***h***^(*N*)^ ∈ ℝ^*M×N*^ to be low-rank. Rank constraint is a popular regularization method (Candes and Recht, 2012) that can improve the model’s robustness to noise in individual neuron spike trains. Therefore, we propose p-RRR, which seeks to maximize

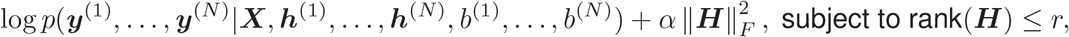

where 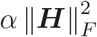 is an additional *L*_2_ regularization penalty to prevent overfitting (Paninski, 2004). *α, r* are hyperparameters which can be determined by cross-validation. *r* is typically taken to be less than *N* (the number of neurons), and when *r* = *N*, p-RRR is equivalent to *N* independent Poisson GLMs. When 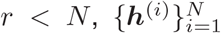 have to be fitted simultaneously in one matrix ***H***. The optimal ***H*** does not have a closed-form solution, but nonetheless can be solved numerically. In practice, we implemented p-RRR as a simple feedforward network in Pytorch (Paszke et al., 2019), and optimized it with the limited-memory Broyden–Fletcher– Goldfarb–Shanno (L-BFGS) algorithm (Liu and Nocedal, 1989).

### Model selection

The two major hyperparameters of p-RRR are the rank constraint (*r*) and the weight for *L*_2_ regularization penalty (*α*). When ***X*** contains multiple types of measurements, since different parts of ***h***^(*i*)^ describe relation hips with different measurements, they are given separate *L*_2_ regularization weights. For example, if 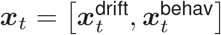, where 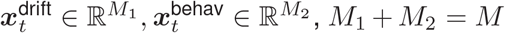, then 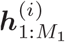 will correspond to 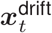, and 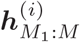 will correspond to 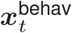. So the *L*_2_ penalty will be 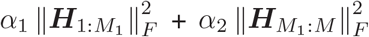, where 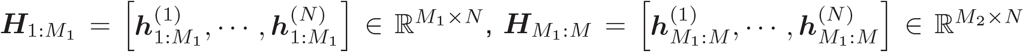, and *α*_1_, *α*_2_ are determined individually.

We used grid search to find the optimal values of *α* and *r*. The typical values we tried for *α* are 0, 10^*−*4^, 10^*−*3^, 10^*−*2^, 10^*−*1^, and for *r* are integers between 1 and 60 (typically as multiples of five). The optimal values of *α* and *r* are determined through a two-fold cross-validation. We split the recorded timestamps (53581 ± 632 per session) into ten even blocks. Alternating blocks were used as training or testing sets. This method was chosen over random sampling to avoid autocorrelations between training and testing data. *α, r* were chosen based on c.v. *D*^2^: we first computed a testing *D*^2^ for each neuron, and then obtained a population average *D*^2^ (weighted by each neuron’s variance on the test set) for each fold. c.v. *D*^2^ is taken as the average population *D*^2^ across two folds. For each *r*, the optimal *α* was chosen to achieve the highest c.v. *D*^2^ across all values of *α*. The optimal *r* was then chosen to be the minimal *r* achieving at least 99% of the maximum c.v. *D*^2^ across all *r*. To identify the rank for 95% performance (Figure 1L), we first created a linear interpolation on the c.v. *D*^2^ from ranks we experimented, and found the minimal rank achieving at least 95% of the maximum c.v. *D*^2^ across all ranks.

### Null predictors

The null term, which accounts for slow changes in electrophysiology (see Figure 1), was computed by low-pass filtering the binned spikes of each neuron with a cutoff frequency of 0.1 Hz. The resulting spike matrix, which includes only the”infraslow” activity, was decomposed using PCA. The first 10 PCA components were used as the null term. This term was included in all models; the Null model only contained this term and an intercept.

### Stimulus predictors

Each unique image was represented by 30 time-shifted binary vectors, each marking one time point relative to stimulus onset with “1” at the corresponding entry (and “0” elsewhere).

### Behavior predictors

For analysis of the face video, motion energy was calculated between frames and then decomposed with SVD using FaceMap (Syeda et al., 2024). For analyses with neural data, motion energy temporal components were linearly down-sampled to the same binning resolution as for electrophysiology and widefield imaging (35 Hz). The behavior predictors consisted of the top 30 temporal components. Each component had 30 time-shifted copies included in the behavior predictor, capturing the dependence on both current behavior and the recent history of behavior.

### Global predictors

For widefield, frames were decomposed into 500 orthogonal spatial and temporal components using SVD. Then, a first-order differencing was applied to the temporal components. The top 30 components and their time-shifted copies (×30) were used. For spiking activity from ACA, neurons with average firing rate below 0.2 spikes/sec were excluded. Other quality control metrics were not applied, as only model targets required stringent identification of single units. Spikes from each neuron were binned at 35 Hz, matching the imaging acquisition rate, and 30 time-shifted copies from each neuron were included in the model.

### Local predictors and targets

After initial quality control metrics (see Electrophysiology), neurons with average firing rate below 0.2 spikes/sec were excluded. These single units were binned at 35 Hz. When used as predictors, 30 time-shifted copies were included per neuron.

### Local model with varying number of peer predictors

See Figure 1K. In each session, 50 V1 neurons were randomly chosen as prediction targets. Each target neuron was predicted with *N* =2, 4, 6, 8, 10, 20, 40, 60, 80, 100, 120, 160, 200, 250, 300 peer neurons (as long as *N* is smaller than the recorded number of neurons) using a single-target prediction model. For each *N, N* peer neurons were randomly selected to form the predictor group, and larger predictor groups were built by adding additional peer neurons to the smaller groups. We repeated this step 10 times for each *N* for each target neuron, and the final result for each target neuron was taken as the average result across 10 repeats. The mean across all target neurons was plotted for each session. Residual Fano factors in Supp Figure 4 were computed similarly, with stimulus kernels included alongside the peer predictors. See section below for residual Fano factor computation. The residual Fano factor for each target neuron was taken as the average residual Fano factor across 10 repeats. The median across all target neurons was plotted for each session; for some sessions individual target neurons are also visualized.

### Known rank peer prediction simulation

See Supp Figure 2. For a neuron population of size *N*, we randomly generated firing rates **Λ** across *T* timesteps as **Λ** = **Λ**_1_**Λ**_2_ + *1:*, where **Λ**_1_ is *T* × *r*, **Λ**_2_ was *r* × *N*, ϵ is Gaussian noise. It follows that **Λ** has a low-rank structure if *r < N*. To match the size of our experimentally recorded spike matrices, we set *T* = 54, 000, *N* = 256. We generated spike trains **Σ** by drawing from Poisson(*αf* (**Λ**)), where *f* is the element-wise softplus function, and *α* is a scaling factor to ensure that **Σ** and recorded spike matrices are of a similar scale. We computed the maximal pairwise Wasserstein distance between the seven recorded spike matrices, and chose *α* such that the Wasserstein distance between at least one recorded spike matrix and **Σ** was less than the maximal pairwise distance. We generated **Σ** with *r* = 2, 4, 8, 16, 32, 64, 128, 256, and repeated the peer prediction analysis (Figure 1J, also see details in section above) on each **Σ**.

### Quantifying explained trial-to-trial variability (the “residual” Fano factor)

Let Y, S, O be random variables representing the binned spike count of some neuron, the stimulus, and non-stimulus experimental measurements, such as other behavioral or neural features. p-RRR is fitted under the assumption Y | S, O ∼ Poisson(*g*(S, O)), where *g* is some function to be modeled. When fitting p-RRR, observations of Y, i.e., *y*_1_,…, *y*_*T*_, are organized into the response vector ***y*** of some neuron; observations of S, O are organized into the design matrix ***X***.

We devised a novel measure, called the residual Fano factor, to quantify the amount of trial-to-trial variability Var(Y|S) explainable by O:

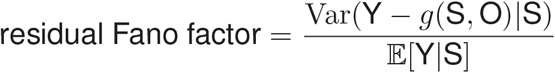

In the following, we show that when the p-RRR assumption Y | S, ∼ O Poisson(*g*(S, O)) is true, the residual Fano factor equals 1.

Rewrite the numerator as

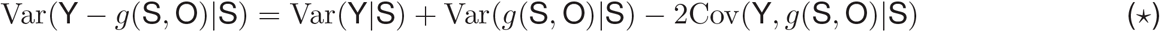

where

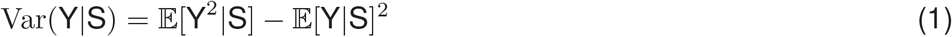

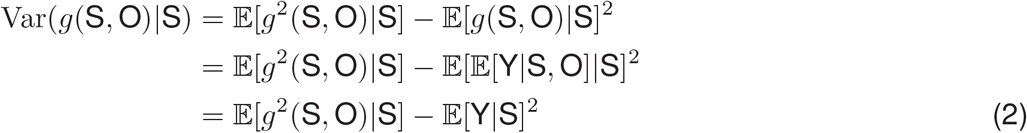

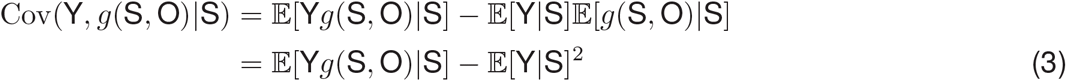

Combine (1)(2)(3),

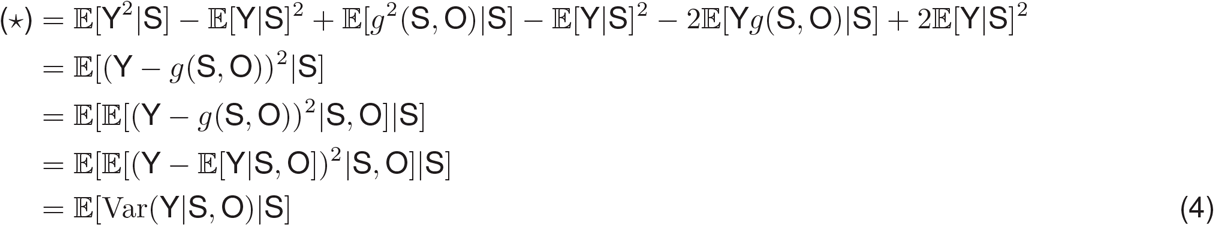

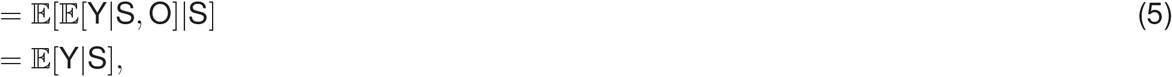

where the equality between (4) and (5) follows from the property of the Poisson distribution. So

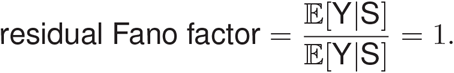

In other words, neural covariates represented by O can account for all trial-to-trial variability down to Poisson noise. If the Poisson assumption does not hold, that is, if Y | S, O ∼ 𝒟 (*g*(S, O)), where 𝒟 is different from the Poisson distribution, then p-RRR may not be the best model for *g*. Nonetheless, if using *g* modeled by p-RRR we still have E[*g*(S, O) |S]= E[Y |S], then we can still follow the calculation to (4). If has sub-Poisson variability, then (4) *<* (5), and residual Fano factor *<* 1. If 𝒟 has super-Poisson variability, then (4) *>* (5), and residual Fano factor *>* 1.

We estimated the residual Fano factor using trial-aligned data and p-RRR prediction on the test set (which was concatenated between the two folds). For each stimulus, to estimate the numerator, we took the difference between the observed spikes and p-RRR prediction, and computed its variance across trials. An estimation of the denominator was obtained by computing the average spikes across trials. The numerators from all stimuli were scattered against the denominators, and the residual Fano factor was taken as the slope *k* of the best-fit linear model *y* = *kx* on these scattered points. In Figure 2, each trial time point had a separate linear fit. In Supp Figure 4, all trial time points were included in a single linear fit.

We also verified the condition E[*g*(S, O) | S] = E[Y | S] for p-RRR models where O is behavior, global brain activity, behavior+global brain activity, or local activity. For every neuron and every stimulus condition, we computed the total observed spikes and p-RRR prediction in the -143ms to 571ms window around stimulus onset for every trial, and tested whether the distributions of observed spikes and p-RRR prediction across trials have the same mean. For all sessions and all choices of O, over 90% of p-RRR predictions across all neurons and all stimulus conditions satisfy the condition (two sample t-test, *α* = 0.01).

### Poisson simulation analysis

For each model, we concatenated the test predictions from both folds to reconstruct the whole time series. For each neuron, the model prediction represented the underlying inhomogeneous rate from which 100 Poisson-random draws were taken. Each draw was scored against the prediction using *D*^2^. In order to compute the z-score, we used the simulated *D*^2^ score distribution as the population, and the prediction *D*^2^ (scored against data) as the raw score.

### Identifying and comparing neural manifolds predicted by p-RRR

We used the “half-half” setup (Supp Figure 2A) to allow for identification of the Local manifold. Half of the neurons in the V1 population were used as predictors and the other half as targets. Stimulus, Behavior, and Global models were also fit to predict the target population. To obtain prediction manifolds, we performed PCA on p-RRR predictions, and took the top PCs accounting for over 99% variance as the predicted manifold.

The alignment between two manifolds is quantified by the amount of activity variance in one manifold captured by the other, with more variance captured indicating a higher degree of alignment. As an example, consider the Stimulus and Behavior manifolds. Take the Behavior model prediction and perform PCA. Suppose the top *R*_Behav_ PCs account for over 99% variance, then the Behavior manifold is *R*_Behav_-dimensional, nd the PCs form an orthonormal basis 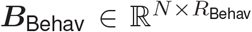 of it, where *N* is the number of V1 neurons. Let **Λ**_Stim_ ∈ R^*T×N*^ be the Stimulus model prediction, where *T* is the number of timestamps. Following Stringer et al. (2019), we can find the dimensions inside Behavior manifold that contain information about Stimulus prediction by identifying a set of ***e***_*i*_ ∈ ℝ^*N*^, *i* = 1,…, *R*_Behav_, such that 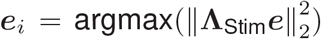, subject to 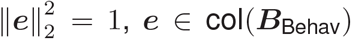, ***e*** ⊥ ***e***_1_,…, ***e***_*i*_ ^−^ _1_. ***e***_*i*_ are given by the columns of ***B***_Behav_***V***, where 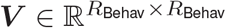contains the right singular vectors of **Λ**_Stim_***B***_Behav_. The percentage of variance on the Stimulus manifold captured by Behavior manifold is 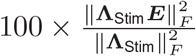, where 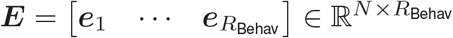

Our manifold alignment metric is closely related to those used in previous studies. Our metric is equivalent to the metric neural variance accounted for (VAF), defined in equation 10 in Gallego et al. (2018). Both metrics share the same denominator, and using the fact that ***E*** has orthonormal columns, we can establish the equivalence of the numerators:

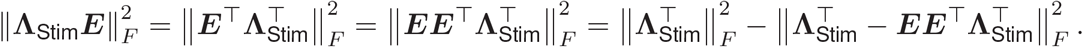

Also, note that 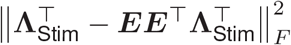 is the squared numerator of the normalized subspace error (*e*_sub_), defined by equation in Gokcen et al. (2022), so our metric also equals to 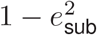.

We call ***E*** the shared dimensions between the Stimulus and Behavior manifolds, with the amount of stimulus-related variance accounted for by individual dimensions decreasing from its first to the last column. The alignment metric is between 0 and 100 (inclusive), where 0 means the Stimulus manifold is orthogonal to the Behavior manifold, and 100 means all modes in the Stimulus manifold can also be found in the Behavior manifold. If the Stimulus manifold has both shared and private modes relative to the Behavior manifold, then the metric is between 0 and 100 (exclusive), with higher values indicating that more stimulus-predicted variance was captured by shared modes.

The reported aligment score was cross-validated using our two-fold cross-validation scheme. Following the previous example, on each fold, we first used **Λ**_Stim_ and ***B***_Behav_ derived from training set predictions of Stimulus and Behavior models to identify ***e***_*i*_. The amount of information captured by each ***e***_*i*_ was then evaluated using **Λ**_Stim_ from the test set. Lastly, the average score of the metric across two folds was reported.

## Supplementary Information

**Supplementary Figure 1:**
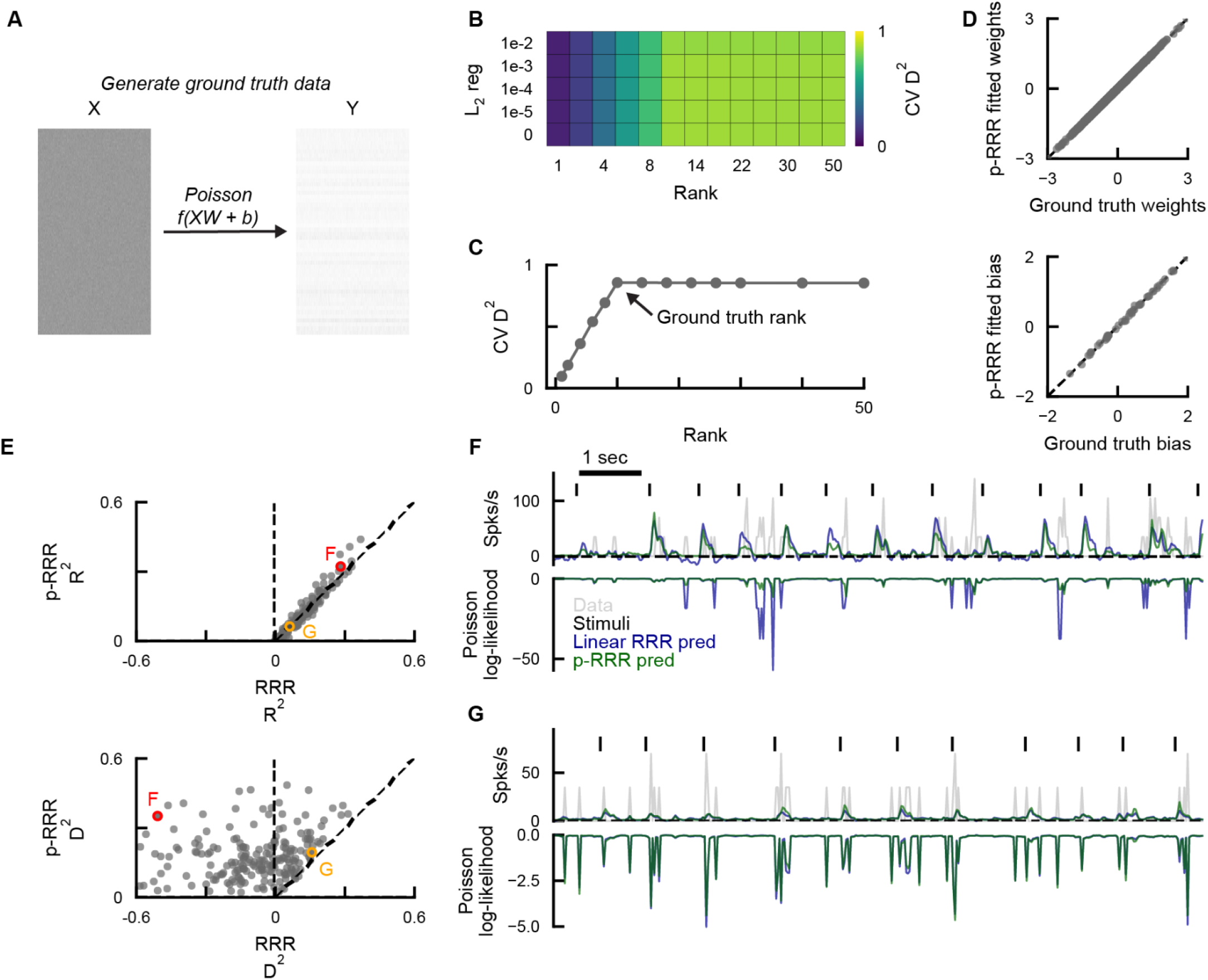
Poisson-RRR recovers ground-truth low-dimensional structure and performs better than linear RRR. **A.** Schematic of ground truth data generation to predict Y from X with a known relationship. We randomly generated X (size 10000 × 100), weights W with a rank of 10 (size 100 × 50), and bias b (size 50). Y (size 10000 × 50) was computed as Poisson draws from f(XW+b), where f was an elementwise softplus nonlinearity. **B.** Cross-validated *D*^2^ performance with varying rank and L2 regularization weight. **C.** Cross-validated *D*^2^ across ranks using optimal regularization per rank. Maximum performance occurred with a rank constraint of 10, consistent with the rank of the ground truth weights. **D.** Top, p-RRR recovered weights versus ground-truth weights. Bottom, p-RRR recovered bias versus ground-truth bias. **E.** Model scores for individual neurons predicted by p-RRR or linear RRR in an example session. Scores are cross-validated R2 (top) or cross-validated *D*^2^ (bottom). Letters indicate example neurons in F and G. **F.** Linear RRR and p-RRR predictions of an example neuron, corresponding to red circles in E. Negative predictions from RRR were set to a small positive number (1e-8) to avoid numerical issues when computing log likelihoods. **G.** As in F, but for the neuron corresponding to orange circles in **E**.

**Supplementary Figure 2:**
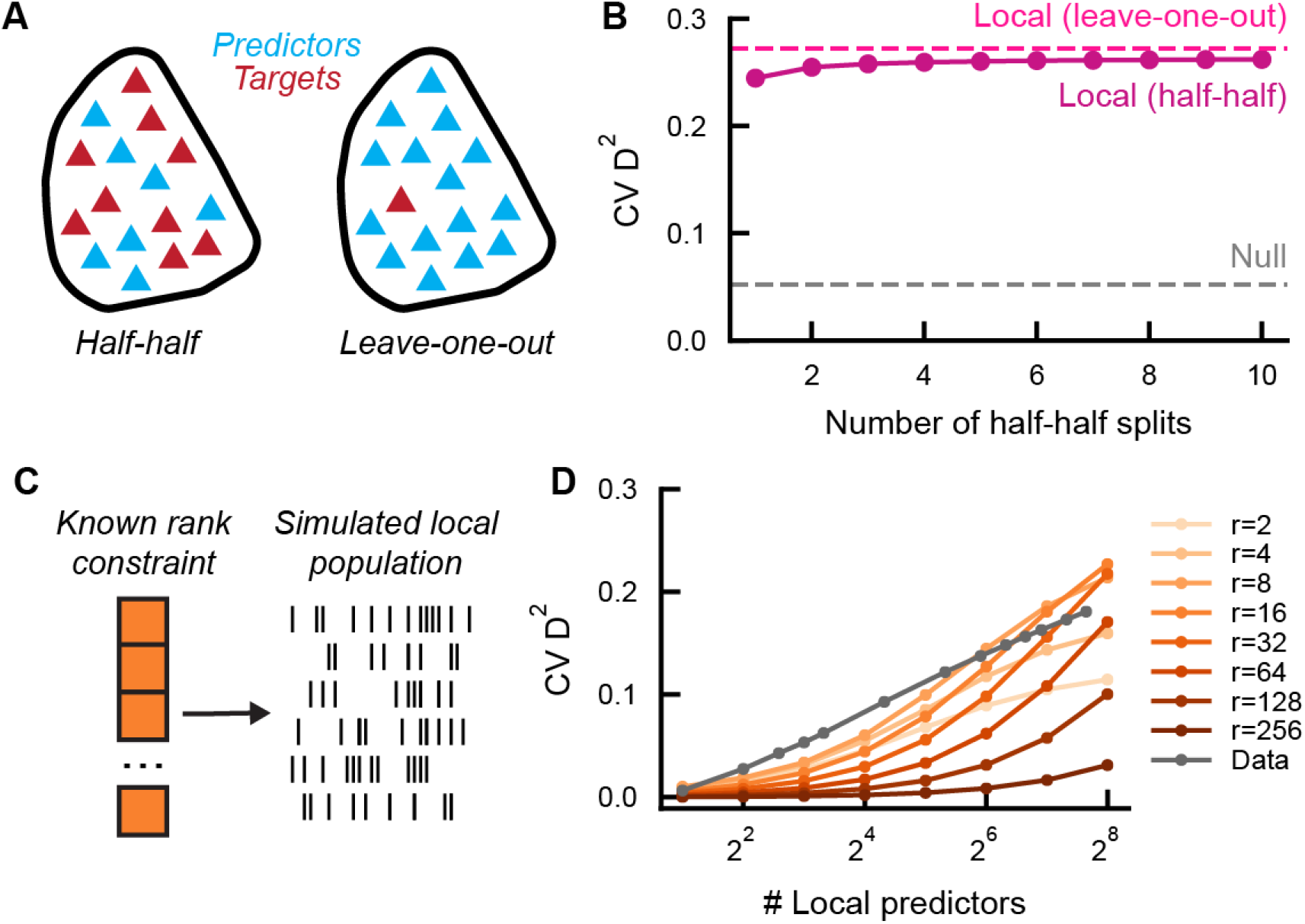
Parameters and rank simulation for the Local model. **A.** Schematic of predictors and targets in variants of the local model. Left, we split recorded VISp neurons into random halves, using one half as predictors and the other half as targets, and fit p-RRR with a rank constraint. This is the setup of the Local model in Figure 1L, M. Right, single-target models were fit for each V1 neuron as an individual target, and with all other simultaneously recorded V1 neurons as predictors. The p-RRR model was equivalent to a Poisson-GLM in this case. This is the setup of the Local model in Figure 1J, K. **B.** Performance of the “half-half” local model as a function of the number of splits. In a single random split, the most predictive peers of some target neuron may be in the targets instead of the predictors, so the prediction of this target neuron may be much worse than the leave-one-out prediction. Averaging across predictions of multiple random splits can mitigate this issue, as seen in the increase in half-half result with increased number of splits. Nonetheless, note that the half-half result with a single split is only slightly worse than the leave-one-out result, suggesting that only a small fraction of V1 neurons are strongly correlated with particular peers, and the majority of them are correlated with shared low-dimensional fluctuations. **C.** Schematic of rank-constrained population simulation. Neurons were randomly generated from a population with a known rank constraint. **D.** Population explained variance (cross-validated D^2^) as a function of the number of local predictors. Colors indicate ranks. Qualitatively, we note that the observed D^2^ increase from experimental recordings (gray line, averaged across sessions) bears more similarity with results on simulated populations with a low-dimensional structure (e.g. r=4).

**Supplementary Figure 3:**
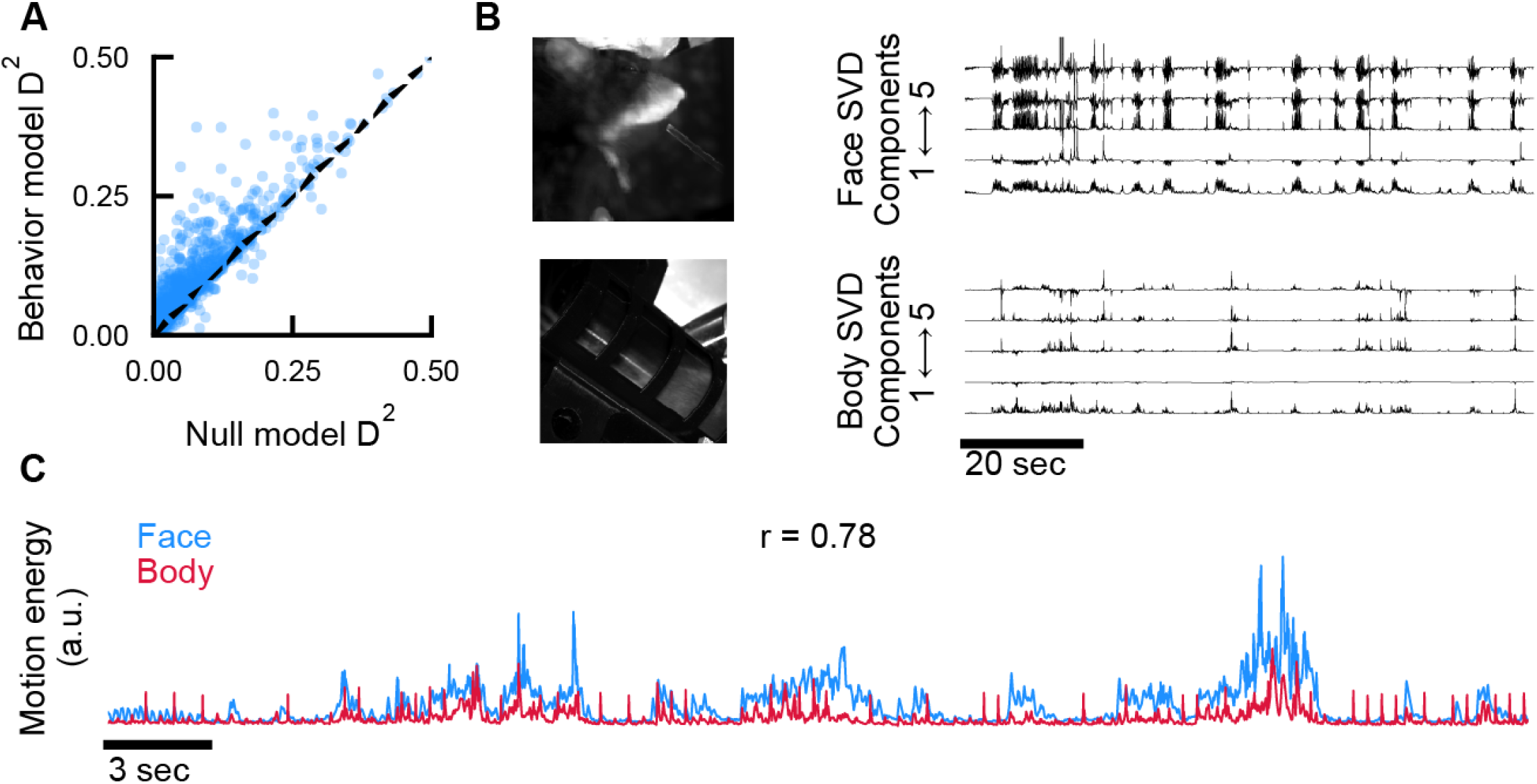
Considerations for the Behavior model. **A.** Cross-validated *D*^2^ scores from Null and Behavior models for individual neurons in all sessions. **B.** Left, example frames from face video camera (top) and body video camera (bottom). Right, example time series of temporal components of SVD from each video. **C.** Overlaid example time series of face and body motion energy. Correlation is computed across the entire session.

**Supplementary Figure 4:**
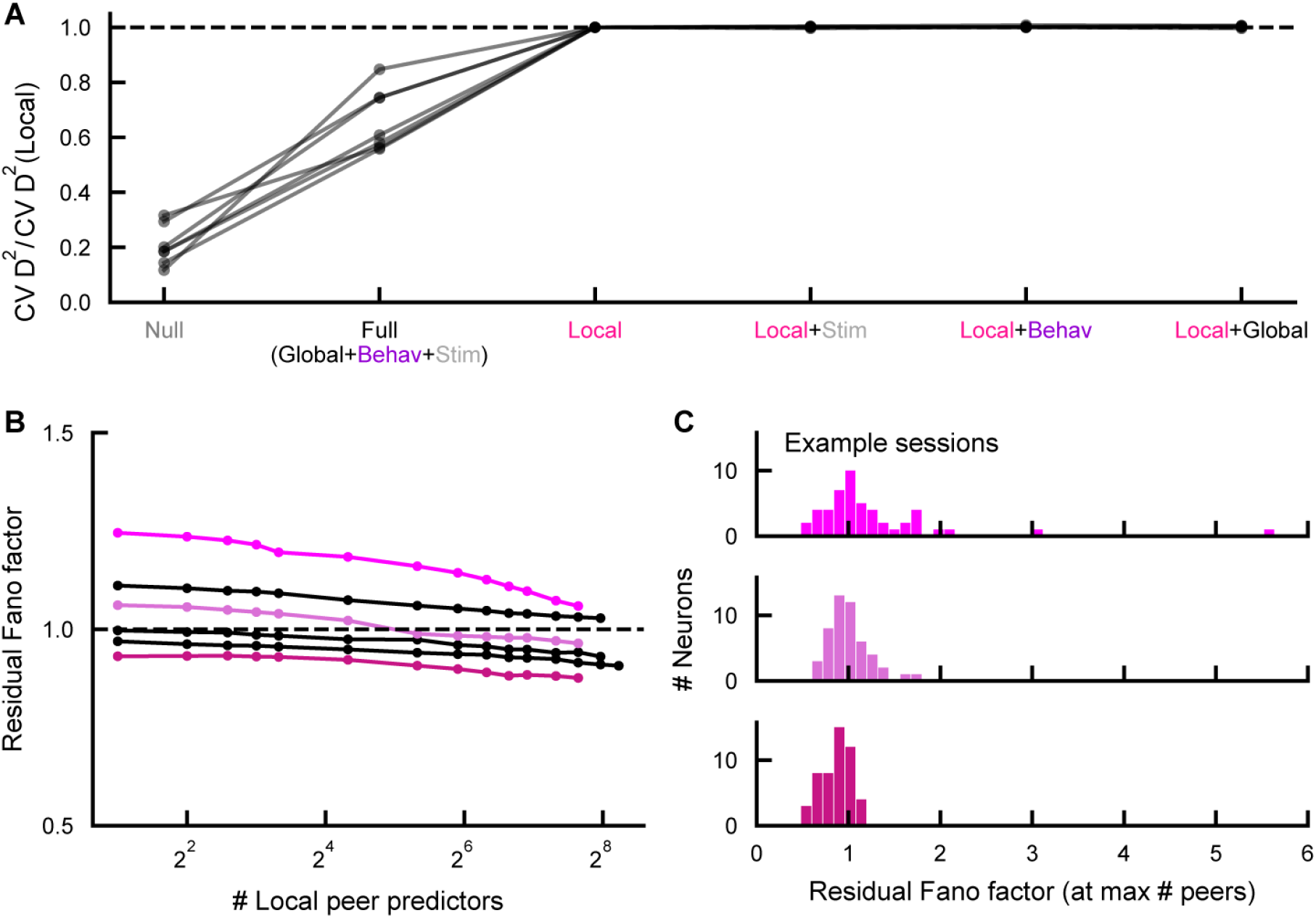
The Local model represents the complete set of shared fluctuations in V1. **A.** Cross-validated deviance explained scores of different models relative to the Local model performance (above 1 indicates better performance than Local model alone). Lines represent sessions. **B.** Residual Fano factor as a function of the number of peer neurons used as predictors in the single-target Local model. Lines represent sessions. Colors indicate corresponding example sessions in **C.** **C.** Distribution of residual Fano factor for individual neurons, after accounting for variance from the Local model. Colors correspond to sessions in **B.**

**Supplementary Figure 5:**
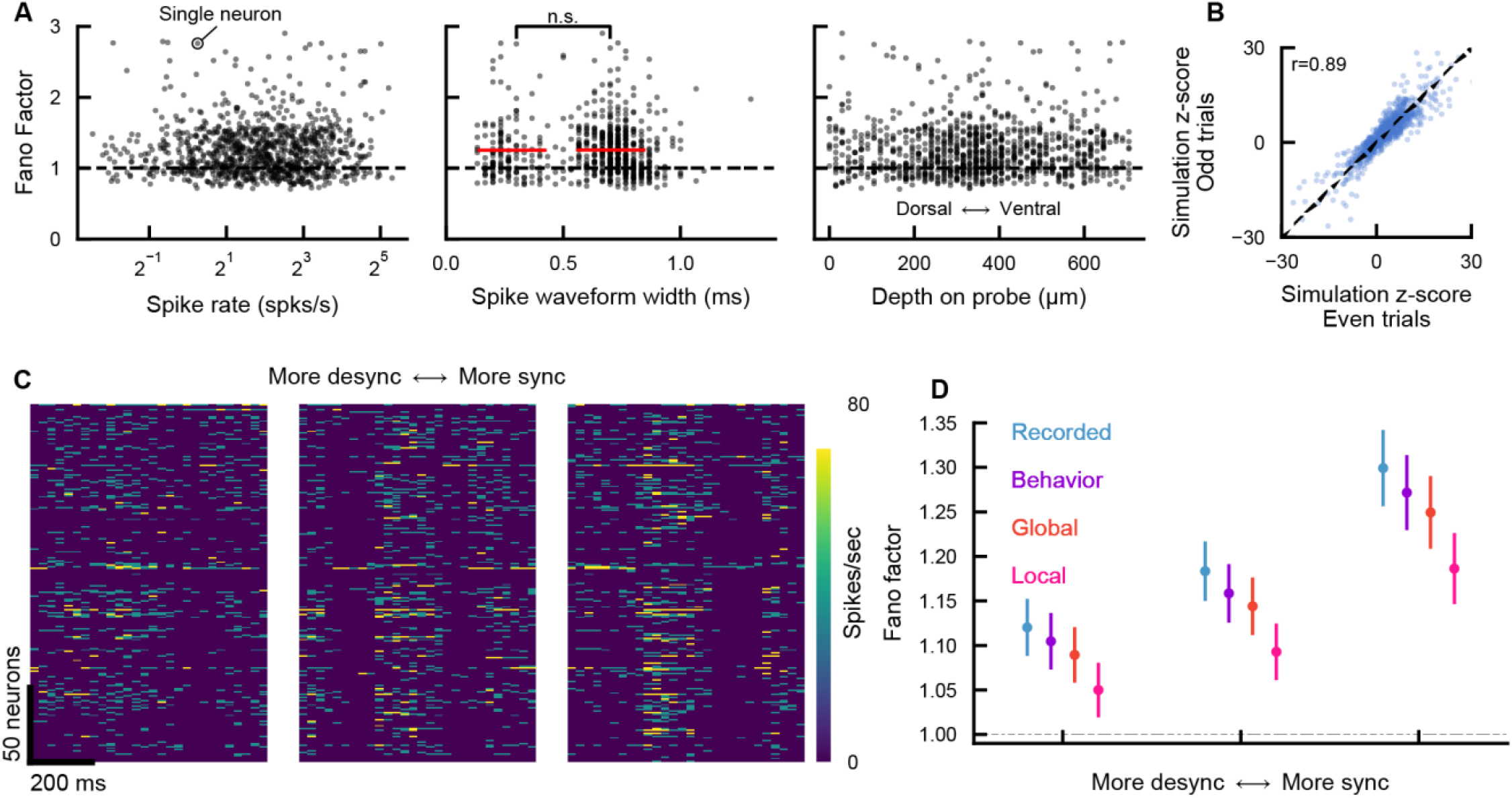
The Poisson measures are reliable, state-dependent, and not correlated with basic neuron properties. **A.** Fano factor as a function of neuron properties. Left, versus spike rate. Middle, versus spike waveform width. Waveform width is computed as the time between peak and trough. Red lines indicate means of narrow and broad waveform neurons (classified by neurons below and above 0.5 ms waveform width, respectively). Right, versus neuron depth on probe. **B.** Poisson simulation z-scores (see Figure 2H-J) computed from even trials (x-axis) and odd trials (y-axis). **C.** Example trials from desychronized and synchronized categories. **D.** Fano factor and residual Fano factor across synchronization categories.

