## Supplementary Figures for "Global and local origins of trial-to-trial spike count variability in visual cortex"

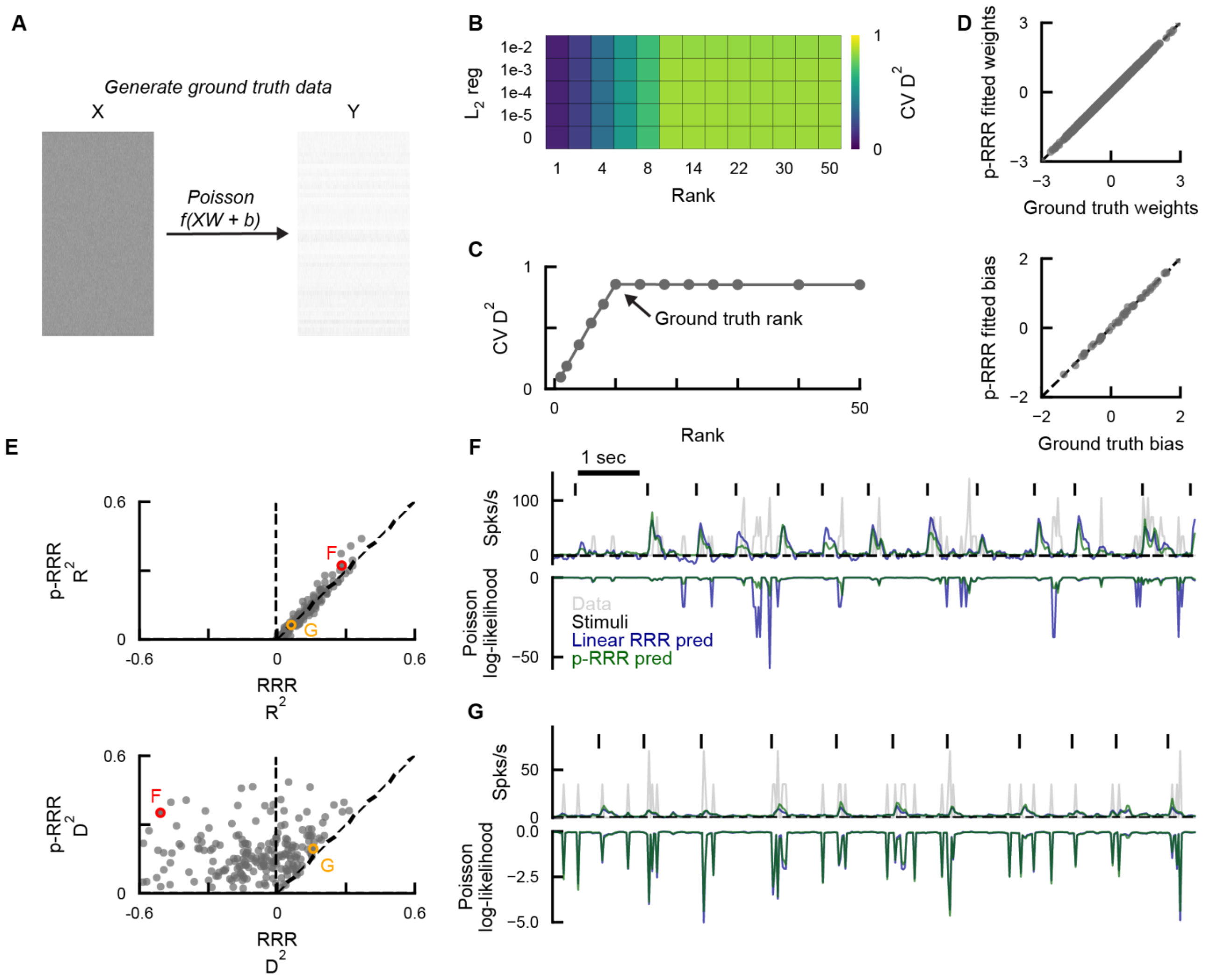


**Supplementary Figure 1: Poisson-RRR recovers ground-truth low-dimensional structure and performs better than linear RRR**

**A.** Schematic of ground truth data generation to predict Y from X with a known relationship. We randomly generated X (size 10000 x 100), weights *W* with a rank of 10 (size 100 x 50), and bias *b* (size 50). Y (size 10000 x 50) was computed as Poisson draws from *f(XW+b)*, where *f* was an elementwise softplus nonlinearity.
**B.** Cross-validated $D^2$ performance with varying rank and $L_2$ regularization weight.
**C.** Cross-validated $D^2$ across ranks using optimal regularization per rank. Maximum performance occurred with a rank constraint of 10, consistent with the rank of the ground truth weights.
**D.** Top, p-RRR recovered weights versus ground-truth weights. Bottom, p-RRR recovered bias versus ground-truth bias.
**E.** Model scores for individual neurons predicted by p-RRR or linear RRR in an example session. Scores are cross-validated $R^2$ (top) or cross-validated $D^2$ (bottom). Letters indicate example neurons in F and G.
**F.** Linear RRR and p-RRR predictions of an example neuron, corresponding to red circles in **E**. Negative predictions from RRR were set to a small positive number (1e-8) to avoid numerical issues when computing log likelihoods.
**G**. As in F, but for the neuron corresponding to orange circles in **E**.

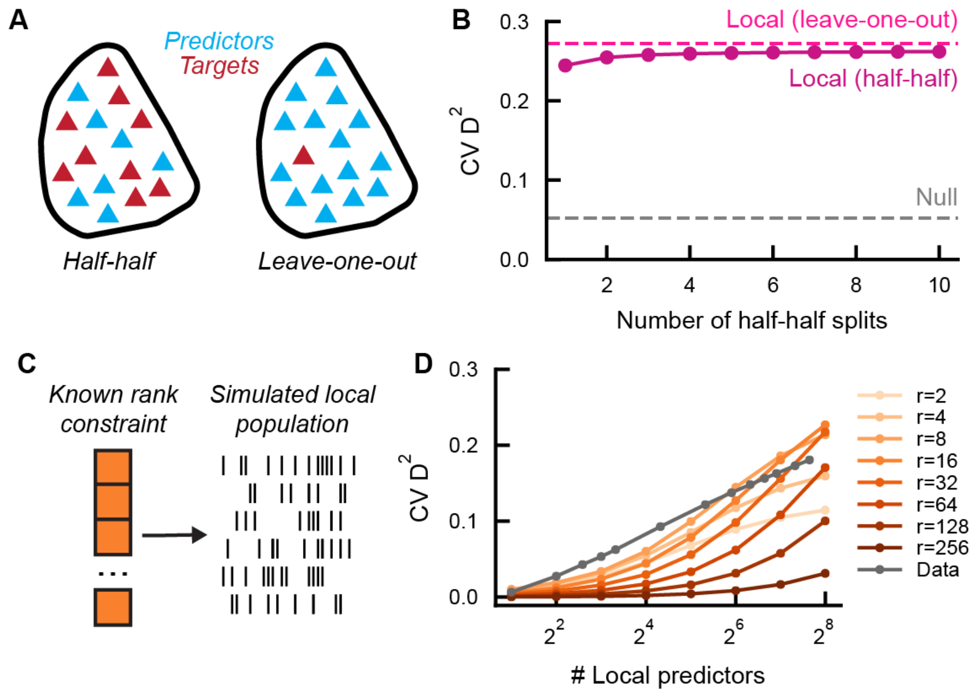


**Supplementary Figure 2: Parameters and rank simulation for the Local model.**
**A.** Schematic of predictors and targets in variants of the local model. Left, we split recorded VISp neurons into random halves, using one half as predictors and the other half as targets, and fit p-RRR with a rank constraint. This is the setup of the Local model in Figure 1L, M. Right, single-target models were fit for each V1 neuron as an individual target, and with all other simultaneously recorded V1 neurons as predictors. The p-RRR model was equivalent to a Poisson-GLM in this case. This is the setup of the Local model in Figure 1J, K.
**B.** Performance of the "half-half" local model as a function of the number of splits. In a single random split, the most predictive peers of some target neuron may be in the targets instead of the predictors, so the prediction of this target neuron may be much worse than the leave-one-out prediction. Averaging across predictions of multiple random splits can mitigate this issue, as seen in the increase in half-half result with increased number of splits. Nonetheless, note that the half-half result with a single split is only slightly worse than the leave-one-out result, suggesting that only a small fraction of V1 neurons are strongly correlated with particular peers, and the majority of them are correlated with shared low-dimensional fluctuations.
**C.** Schematic of rank-constrained population simulation. Neurons were randomly generated from a population with a known rank constraint.
**D.** Population explained variance (cross-validated $D^2$) as a function of the number of local predictors. Colors indicate ranks. Qualitatively, we note that the observed $D^2$ increase from experimental recordings (gray line, averaged across sessions) bears more similarity with results on simulated populations with a low-dimensional structure (e.g. r=4).

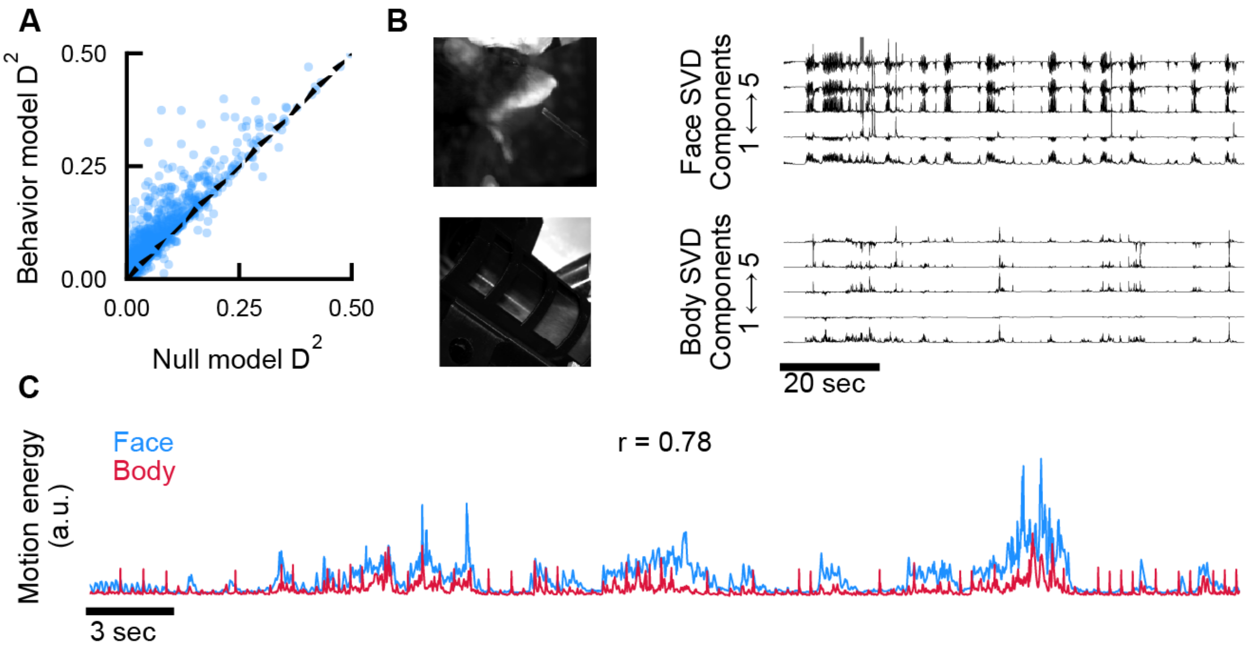


**Supplementary Figure 3: Considerations for the Behavior model.**

**A.** Cross-validated $D^2$ scores from Null and Behavior models for individual neurons in all sessions.

**B.** Left, example frames from face video camera (top) and body video camera (bottom). Right, example time series of temporal components of SVD from each video.

**C.** Overlaid example time series of face and body motion energy. Correlation is computed across the entire session.

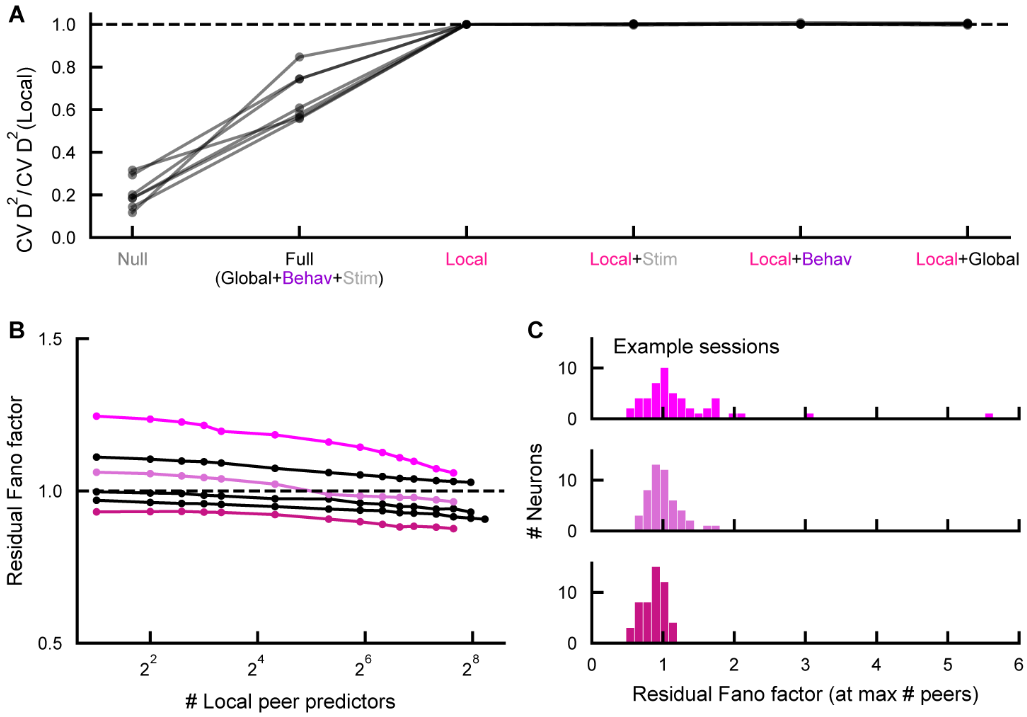


**Supplementary Figure 4: The Local model represents the complete set of shared fluctuations in V1.**

**A.** Cross-validated deviance explained scores of different models relative to the Local model performance (above 1 indicates better performance than Local model alone). Lines represent sessions.

**B.** Residual Fano factor as a function of the number of peer neurons used as predictors in the single-target Local model. Lines represent sessions. Colors indicate corresponding example sessions in **C.**

**C.** Distribution of residual Fano factor for individual neurons, after accounting for variance from the Local model. Colors correspond to sessions in **B.**

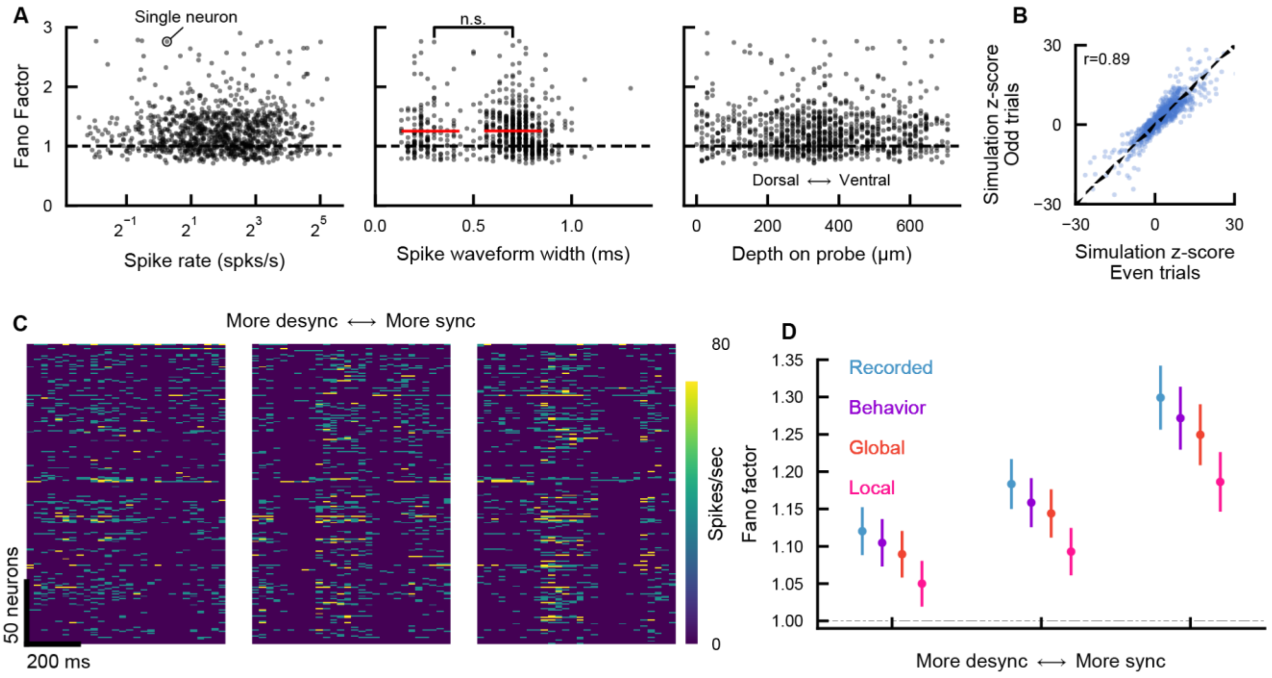


**Supplementary Figure 5: The Poisson measures are reliable, state-dependent, and not correlated with basic neuron properties.**

**A.** Fano factor as a function of neuron properties. Left, versus spike rate. Middle, versus spike waveform width. Waveform width is computed as the time between peak and trough. Red lines indicate means of narrow and broad waveform neurons (classified by neurons below and above 0.5 ms waveform width, respectively). Right, versus neuron depth on probe.

**B.** Poisson simulation z-scores (see Figure 2H-J) computed from even trials (x-axis) and odd trials (y-axis).

**C.** Example trials from desychronized and synchronized categories.

**D.** Fano factor and residual Fano factor across synchronization categories.
